# Walking in circles: Linking high- and low-level parameter scaling of visually guided and spontaneous turning behaviour

**DOI:** 10.64898/2026.07.01.735770

**Authors:** Merit Meschenmoser, Volker Dürr

## Abstract

The ability to change one’s current heading, i.e. to turn, is essential for all walking animals. While several studies have addressed how leg movement or inter-leg coordination may change during turning, relatively little is known about how turning-related changes scale with turn magnitude. Here, we used spontaneous and visually induced turns of unrestrained walking stick insects to test (i) how high-level parameters of unrestrained turning scale with low-level parameters of leg movement, and (ii) the effect of visual guidance on turning parameters. To this end, we used a step change in stationary landmark position in an open-field arena to constrain timing and magnitude of target-directed turns. These visually guided turns were compared with spontaneous turns in an all-white condition. We show that visually induced turns were walked at a larger forward velocity and had fewer short steps than spontaneous turns. The scaling of turning responses was dominated by an increase in turning duration (factor 1.87) rather than turning speed (factor 1.32). Increased rotational velocity correlated with reduced forward velocity, though with flexible timing of both effects. These changes were accompanied by larger shifts in step direction, as well as an increased asymmetry of step types between inner and outer legs, suggesting a mix of distinct turning strategies, that depend on overall turn angle. Future models on six-legged locomotion should thus consider the incorporation of more than one mechanism to govern turning.

**Summary statement:** Freely-walking stick insects turn by a combination of scaled changes in step-parameters and frequency of distinct step-types.

## Introduction

In terrestrial locomotion, complex movement sequences as part of searching, climbing or turning behaviours, require not only appropriate temporal coordination of several legs, but also adaptation of spatial step parameters such as step length and direction (Ritzmann and Büschges 2007; Cruse et al. 2009a; Büschges and Ache 2025). With regard to turning and curve walking, established knowledge about “low level” step parameter changes in stick insects (Dürr and Ebeling 2005; Cruse et al. 2009b; Gruhn et al. 2009), cockroaches (Franklin et al. 1981; Nye and Ritzmann 1992; Jindrich and Full 1999; Ridgel et al. 2007) and other insects (e.g., Strauss and Heisenberg 1990; Frantsevich and Cruse 2005; Zolotov et al. 1975) has been complemented by insights into the “high level” descending control of turning trajectories (Yang et al., 2024; Rayshubskiy et al., 2025; Cruz and Chiappe, 2023). Still, the direct link between low-level changes in leg movement and high-level changes in body trajectory is not well understood, particularly not in unrestrained locomotion.

Previously, turning in stick insects has been elicited visually through large-field motion cues (Dürr and Ebeling 2005; Gruhn et al. 2009) or salient, stationary landmarks (Kalmus 1937; Rosano and Webb 2007; Zeng et al. 2020; Meschenmoser and Dürr 2025). However, with either stimulus type, it is unknown how turning scales with stimulus magnitude. For example, in order to cover twice the turn angle, animals could either turn for twice as long or twice as fast, or with a combination of both. Our experiments were designed to test for these alternatives, employing an instantaneous displacement of a visual target to elicit two different turn angles within a defined region of our setup.

While it is known that changes in leg movement parameters during turning can be related to the curvature of the walked path (e.g., Jander 1985), adaptive control of (at least) 18 degrees of freedom allows insects to employ various strategies to walk the same curve. Nevertheless, current locomotion models typically implement only a single mechanism that governs turning. Recently, individual idiosyncrasies have been shown to shape straight walking behaviour in *Drosophila melanogaster* (Godesberg et al. 2024). However, this description of inter-individual variation is not yet fully integrated in an understanding of inter-trial variation, which is necessary to tell adaptive control of speed and heading from redundancy-related variability of movement. Here, we test whether insects constrain this redundancy in a context-dependent manner by comparing visually guided with spontaneous turning.

Although stick insects have comparatively unspecialized legs, turning is thought to be initiated by strong changes in front leg stepping (Dürr and Ebeling 2005) and reduced coupling to ipsilateral middle and rear legs (Dürr 2005). Accordingly, front legs have been proposed to be the primary target of visual descending information about how to change heading. Indeed, simulations involving active changes in front leg movement only, with middle and hind legs following passively, have reproduced similar turns as performed by stick insects (Rosano and Webb 2007). More generally, front legs show the largest degree of motor flexibility of all leg pairs, as reflected by their key function in near-range searching (Dürr 2001; Berg et al. 2013; Dürr and Mesanovic 2023), local probing (Bläsing and Cruse 2004a) and targeted reaching (Schütz and Dürr 2011). When studying step cycle parameter variation during unrestrained walking and climbing, Theunissen and Dürr (2013) found that step length distributions of all leg pairs were bimodal. Given the distinct properties of step cycles sampled from these two modes, they suggested that insects generate two distinct classes of step cycles: While “long steps” were proposed to serve propulsion, “short steps” were interpreted as correction steps in case of insufficient foothold. As shorts steps occurred more frequently on edges of objects during climbing, they could also serve repetitive probing or exploration, as proposed for short steps before and during gap-crossing maneuvers (Bläsing and Cruse 2004a). Assuming an exploratory function of intermittent short steps, we expect the frequency of short steps to increase as spatial information about the environment becomes less reliable. Accordingly, we test the hypothesis that the absence visual orientation cues lead to increased exploratory effort, i.e. more short steps, to compensate for the lack of visual spatial cues.

Our experiments thus test the effect of visual guidance on unrestrained turning responses, along with the effect of stimulus magnitude, i.e. the step change in target position. To this end, we used two displacement magnitudes (60°, 120°) and an all-white stimulus to compare visually guided and spontaneous turns of equal curvature. A stationary landmark with no displacement was included as a control for ‘straight walking’. Our analysis links two levels of description of turning behavior: First, we analyze body axis kinematics, relating timing and magnitude of forward translational and rotational velocities; Second, we relate these changes to step cycle parameters of the front legs, with a focus on step length and swing direction.

## Material and Methods

### Experimental animals and setup

We collected data from 52 adult, female stick insects of the species *Carausius morosus* (Sinéty 1901). Of these, 20 were use in pre-experiments and 32 in the main experiment. Specimens were taken from a parthenogenetic breeding colony at Bielefeld University. The animals were kept at a 12h:12h light/dark cycle and fed ad libitum with blackberry leaves and chinese cabbage.

Animals were placed manually into a circular open-field arena with a diameter of 120 cm, surrounded by a translucent screen of 20 cm height. Using external projectors, the screen was illuminated with white light from the outside. In most conditions, a black bar was projected onto the white screen, serving as a visual landmark. The bar was 20° wide as seen from the arena center. Animals were free to walk on the black wooden arena floor. Top-view videos were collected at a frame rate of 50 Hz, using a custom-built camera gantry that was equipped with an infrared-sensitive digital video camera (Basler A602f-2 or A602fc-2, image size: 640×480 px) and a manual zoom lens (Computar, M6Z1212-3S). For a detailed setup description see Niemeier et al. (2021) and Meschenmoser and Dürr (2025).

### Experimental Procedure

In trials with a landmark, animals were always placed in the arena opposite to and facing the landmark. To induce a sudden shift of landmark position, the experimenter manually triggered a step change in landmark position when an animal crossed the midline of the arena, using a wireless remote control (Kensington K72426EU). The magnitude of this landmark displacement varied among conditions and was set by a custom-written MATLAB script. In ‘all-white’ control trials without a landmark, animals were put in the arena opposite to where the black bar would have been in landmark trials (Fig 1 A). The person handling the animal was unaware of the order in which conditions were presented. Time stamps of the change in landmark position were saved in a text file. The video recording was terminated once an animal reached the arena wall or stopped walking.

**Figure 1:**
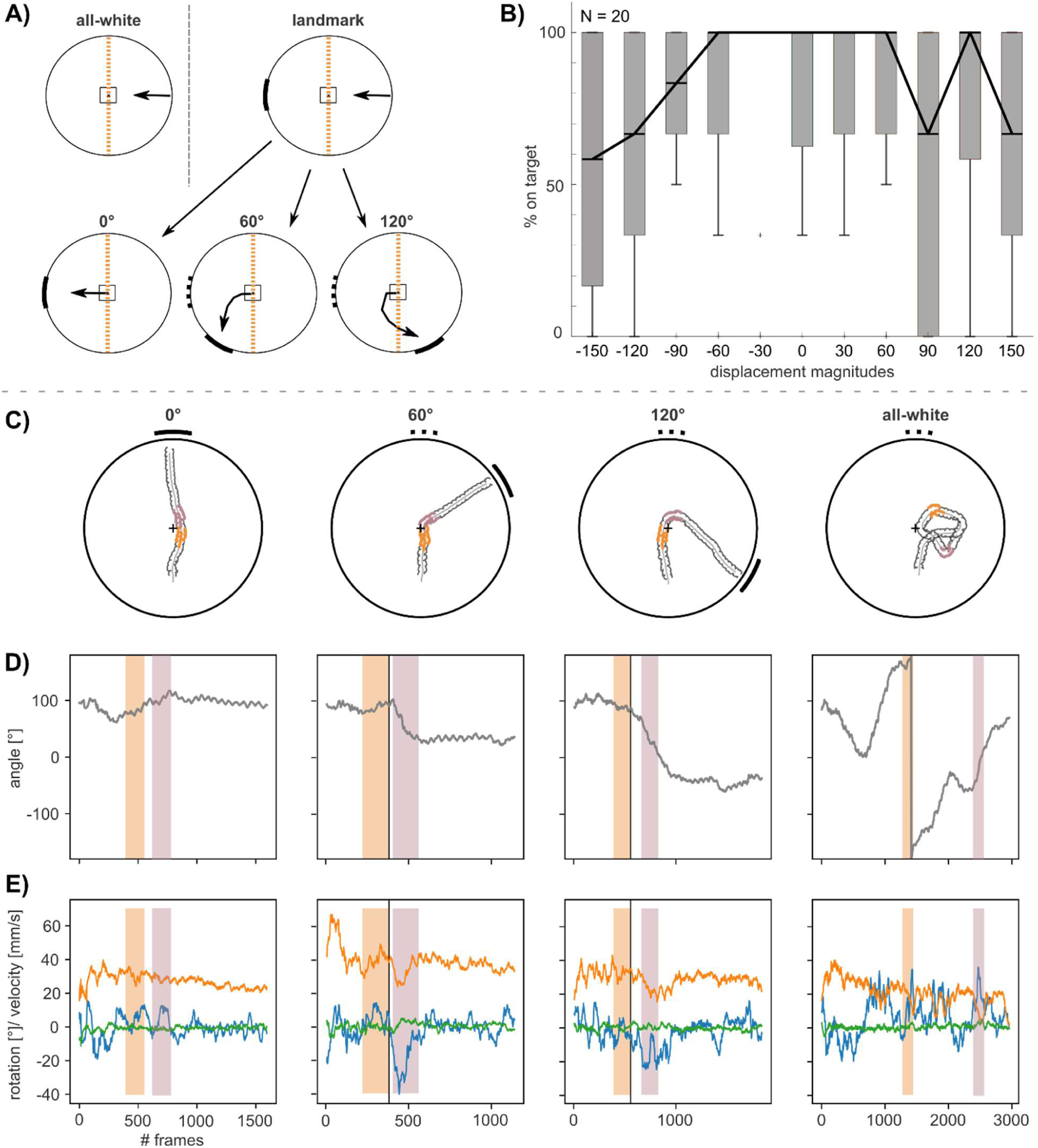
Experimental design and example trials. **(A)** Schematic drawing of the circular arena and experimental procedure. In landmark trials, the position of the landmark was changed as soon as the animal reached the arena center. All-white trials served as controls. Solid black lines outside the arena represent the landmark position and width. Dotted black bars in the second row show the original position of the landmark before the displacement. The black arrow in the upper row shows the start orientation of the animal. Arrows in the second row show the hypothesized orientation response of the animal. The black cross and the dashed orange line highlight the center point and midline of the arena, respectively. **(B)** Boxplots show the likelihood of an animal to be ‘on target’ for different displacement magnitudes, with N = 20 means per animal, from 3 trials per animal and displacement. **(C-E)** Random example trials from the same animal. Suppl. movies 1-4. **(C)** Top view of circular arena. The light grey trace shows the trajectory of the metathorax marker, while the dark grey lines are the trajectories of the tarsal tips of both front legs. Bold, colored parts of the trajectories highlight the two episodes A and B. **(D)** Body orientation throughout a trial. The vertical black lines indicate when the landmark was displaced in 60° and 120° trials. Body orientation angles above or below ±180° were converted to angles within the ±180° range. Colored shading marks the two episodes A and B. **(E)** Three movement components of the center of mass on a planar surface: rotational (blue), forward (orange) and sideward translational (green) velocities. Colored shading marks the two episodes A and B.

Trials were scored as ‘at wall’ and ‘on target’ trials. An animal was considered ‘at wall’ if its head was less than 7 cm away from the wall. To be ‘on target’ animals had to be ‘at wall’ in the target range. This was defined as the target width plus a tolerance of five degree on either side of the target. As there was no target present in the all-white condition, trials could be ‘at wall’ but not ‘on target’.

#### Pre-experiment

A pre-experiment was carried out to determine the reliability of target approach for different magnitudes of landmark displacement. For each trial, we noted whether or not the animal reached the target. No videos were recorded. Our aim was to decide on two magnitudes which were considerably different while guaranteeing high likelihoods of target approach. To this end, we used ten different displacement magnitudes (30°, 60°, 90°, 120°, 150°) and tested them in both clockwise and anti-clockwise directions. Trials with a stationary landmark (0°) were used as a control. We used 20 animals and repeated each condition three times per animal in a pseudo-randomized manner, resulting in 33 single trials per animal. The results showed that the median likelihood to be ‘on target’ decreased at displacement angles beyond ±60°, but was never less than 60% (Fig. 1 B).

#### Experiment

Based on the results of our pre-experiment, we decided on ±60° and ±120° as displacement magnitudes (Fig. 1, A). This choice covered a large portion of the visual field and allowed for simple assessment of linear scaling of response parameters with displacement magnitude.

Data was collected in sets of five trials consisting of one 0° trial, two 60° trials and two 120° trials (one for each direction). The order in which the conditions were presented was randomized within each set. The starting orientation (90° or 180°) of the trial was randomized across all trials, including all-white trials, to exclude confounding, setup-specific effects. Experiments were done in two cohorts which differed slightly in the number of all-white trials and the number of sets. The first cohort experienced one all-white trial after two sets, whereas the second cohort experienced one all-white trial after every set. Additionally, we decreased the number of sets from 8 in the first cohort to 7 in the second cohort. This resulted in 44 trials per animal for the first cohort and 42 trials for the second cohort, respectively. In total we collected 1252 trials from 32 animals. One animal was excluded from the analysis as it only contributed two at-wall trials to the dataset. The number of animal means used in the analysis varied between 26 and 31, as not each animal contributed data points for all response categories. Figure captions state the number of animals included.

### Marker-less tracking and categorization of trials and episodes

Data processing and analysis was done using Python 3.9.13 (Van Rossum et al. 2009) and MATLAB 2022b (The MathWorks Inc. 2022). ChatGPT 4o mini (OpenAI) was used to aid the development of code for data analysis.

Digital videos were analyzed so as to reconstruct body and leg positions, body axis orientation and head position. We used DeepLabCut (Mathis et al. 2018, Nath et al. 2019, Arent et al. 2021) to track several features on the body and the legs of the animal. Tracked features along the body axis were the midpoint between both hindlegs on the metathorax (T3), the midpoint between both middle legs on the mesothorax (T2), the midpoint between both forelegs on the prothorax (T1) and the tip of the head (Hd). Additionally, we tracked four features per leg: the coxa-trochanter joint (CTr), the femur-tibia joint (FTi), the tibia-tarsus joint (TiTa) and the distal tip of the tarsus (Tars) (see Suppl. Fig. 3). Only frames, with a mean confidence rating above 95% across all features were used for further analysis. A median filter (width = 3 frames) and a Gaussian kernel (full width at half max = 5 frames; σ = 2.12, kernel width = 15 frames) were used to smooth feature trajectories. The body orientation was determined as the angle between the vectors T3→T1 and (0, 1)^T^ (see Fig. 2, C).

**Figure 2:**
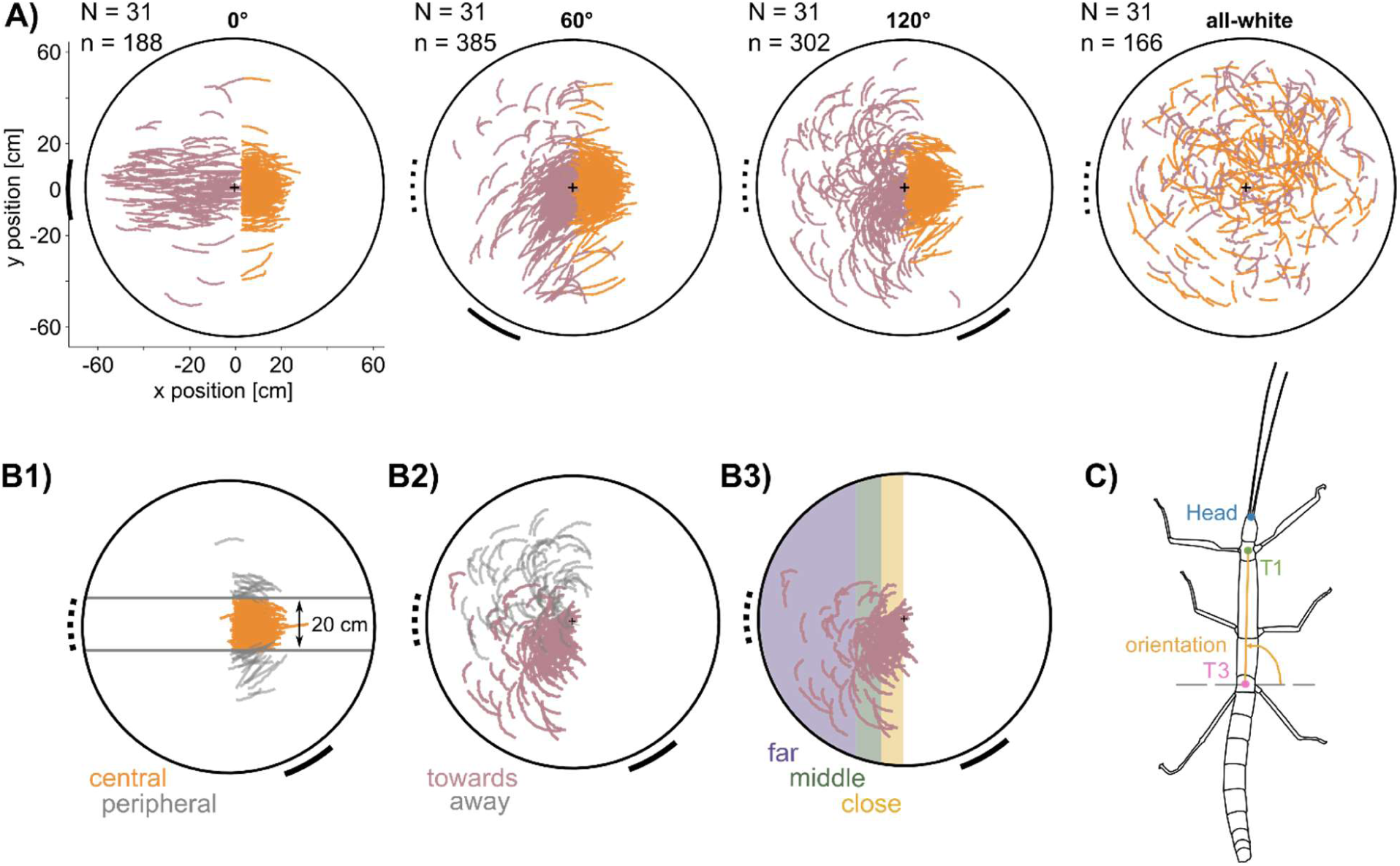
Episode overview and categorization of turning responses. **(A)** Top view of circular arena. Solid black lines outside the circle show the width and position of the black bar presented. Dotted black lines show the first position of the black bar before it was displaced in 60° and 120° trials. In the all-white trial, the dotted line indicates the start orientation of the animal, since no black bar was present at any time. The black cross indicates the midpoint of the arena. Circular plots show the prothorax (T1) positions during two episodes of each ‘on-target’ trial. For the all-white condition, all trials are shown. The two episodes are: A) pre-turning (orange) and B) turning (soft pink). The number of plotted trials is shown on the upper left. Note that the spread of all trajectories does not reflect the likelihood of animal position in the arena, as it strongly overrates peripheral trajectories (see Fig. 4A). **(B1-3)** Schematic drawing of the categorization process of turning responses according to their spatial occurrence. **(B1)** Only ‘central’ trials in which T1 stayed within a corridor of ±10 cm relative to the midline at the end of episode A (orange) were used for further categorization. **(B2)** Depending on turning direction during episode B (pink), trials were categorized as ‘away’ or ‘towards’. **(B3)** ‘Towards’ trials were further categorized as close (yellow), middle (green) and far (purple) turns, depending on the location of episode B (pink). **(C)** Schematic drawing of a stick insect. Colored dots show the thorax features that were used for calculating the body axis orientation of an animal.

A trial started when the animal no longer had contact with the experimenters’ hand and ended either upon reaching the arena wall or if the animal stopped. Stopping was defined as the instant when forward velocity dropped below 1.0 mm/s for at least 1.5 seconds. For each trial, two episodes were compared, each one lasting three seconds. Episode A started three seconds before the change in landmark position. In 0° control trials, it started three seconds before the animal crossed the midline of the arena. In all-white trials, it started at half of the trial. Episode B comprised the three seconds interval with the maximum mean rotational velocity across (Fig. 2 A).

Only ‘*on target*’ trials were used for further analysis of landmark trials. The number of animals varied for certain parts of the analysis due to the fact that not all animals were ‘*on target*’ in all conditions (as indicated in respective figures). All quantitative analyses and statistics were carried out on per-animal means. If not stated otherwise, we used a Friedman-Test and post-hoc Wilcoxon signed-rank tests for matched pairs for statistical comparisons.

### Data analysis at the trajectory level

Kernel density estimates were calculated using the *gaussian_kde* function from *scipy.stats* with a 120cm×120cm grid and a grid resolution of 1cm×1cm. To allow for a comparison between different conditions, density estimates were normalized by the maximum density across all stimuli.

The overall curvature (or tortuosity) of a trial was calculated as the walked distance divided by the length of the shortest possible path, i.e. a straight line. Rotational velocities were calculated for ‘*on target’-*trials only, using the filtered T1 and T3 positions (Sawitzky-Golay filter: polynomial order 1, window size 25 frames). Since trajectories varied strongly, particularly in 120° trials, we categorized turning responses according to the location and direction of maximum turning to facilitate comparison between conditions (Fig. 2B): To exclude ‘peripheral’ trials in which animals left a straight corridor before landmark displacement, only ‘central’ trials in which the prothorax, T1, remained within 10 cm to either side of the midline at the end of episode A were used for further categorization (Fig. 2B1). To account for turning direction, all trials in which the direction of rotation during episode B was directed away from the second landmark position were categorized as ‘away’ turns and not considered further (Fig. 2B2). To account for latency of turning, trials were further categorized according to the location of T1 at the start of episode B. If T1 remained within 10 cm from the arena center, the trial was categorized as a ‘close’ turn. Between 10 and 20 cm from the arena center, it was categorized as a ‘middle’ turn, and everything beyond 20 cm was considered a ‘far’ turn (Fig. 2B3). For the comparison of maximum rotational velocity, timing of maximum rotational velocity and duration of turning, we used ‘close turns’ only. Since forward velocities decreased towards the end of a trial, minimum forward velocities were calculated for the turning episode of close turns only. For this, we used the bootstrapped rotational velocities. The window that we used for the minimum forward velocities, started at the time of the landmark displacement up to the point when the rotational velocity finally fell within the mean plus three times the standard deviation of the rotational velocity before the landmark displacement again (see also Fig. 5C).

For the segmentation of trials into curved and straight parts, trajectories of the two thorax features T1 and T3 were strongly smoothed, using a Butterworth filter (4^th^ order, cutoff frequency: 0.25 Hz), and rotational velocities were recalculated using the filtered T1 and T3 positions. We then used the free library *Ruptures* from python (Truong et al. (2020)) to segment parts of trajectories based on their rotational velocities. To this end, we chose a non-parametric rank model which was robust against outliers. The minimum segment size was set to 150 frames (3 s) and was controlled by a penalty parameter which was set to 50. 389 out of 4940 segments (7.9 %) were shorter than 150 frames but all segments were larger than 100 frames. Performance of the splitting process was evaluated by skimming through randomly chosen trials and visually assessing whether the splitting algorithm performed as intended by looking at T1 trajectories (see Suppl. Fig. 2).

Each trajectory split was resampled to contain 400 points, using linear interpolation between frames. The curvature of a given split was calculated using the metathorax marker T3 and the 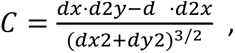, where dx and dy are the first derivatives, while d2y and d2x are the second derivatives of x and y, respectively. Afterwards, the radius of curvature of a segment (ROC) was calculated as 1/median(C).

To compare the forward velocity of similarly curved segments with and without visual guidance, we used the *curve_fit* function from the SciPy *Optimize* package (Virtanen et al. (2020)) to fit the saturation curve

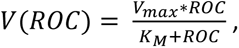

where V_max_ is the maximum forward velocity and K_M_ is the ROC at half V_max_. This was done separately for each animal. To do so, right and left turns were pooled. The prior estimates for V_max_ and K_M_ were 40 and 100 respectively. We used Wilcoxon’s signed rank test for matched samples to compare V_max_ and K_M_ between the all-white and landmark conditions.

### Step cycle detection

We trained six random forest classifiers (scikit-learn, Pedregosa et al. (2011)), one for each leg, to determine whether or not a leg had ground contact. We used 1353 features to train each classifier. The feature vector contained the angles between each leg segment (using their 2D projections), the velocity of three tracked leg features (femur-tibia joint, tibia-tarsus joint and the tarsal tip) as well as the length of three leg segment projections (femur, tibia and tarsus) for all six legs (for details see Suppl. Fig. 3). In addition, we used the instantaneous body rotation, forward and translational velocity as features. All features connected to the legs were included as time series data from 25 frames. For each frame, the feature vector therefore contained information about leg posture and velocity in 12 preceding and 12 subsequent frames. The training set to train the classifiers contained 14916 manually labeled frames, taken from two random trials per animal. From each trial, two random one-second sequences were used. For each frame, the state of each leg was noted as 1 if having “no ground contact”, e.g. during swing, or as 0 if having “ground contact”. The evaluation of the six classifiers showed only minor differences in accuracy (true positives and true negatives) between the three leg pairs (Table 1). Overall, the accuracy of all classifiers was above 95% for front legs and above 97% for middle and hind legs. For the decision “no ground contact” each classifier was more likely to generate false negatives (e.g., stance instead of swing) than false positives (e.g., swing instead of stance), suggesting that leg postures without ground contact were harder to detect as such than leg postures with ground contact.

**Table 1:**
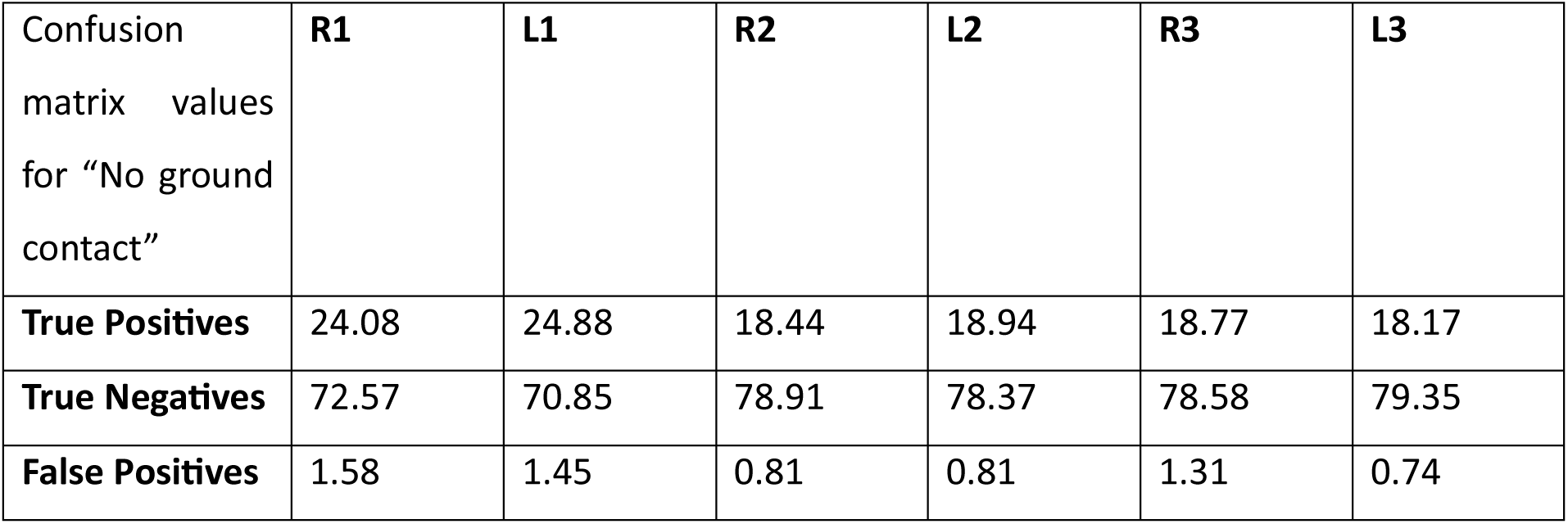
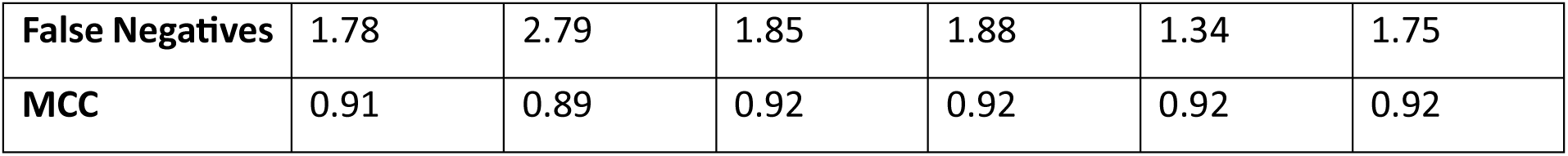
Evaluation results of the six individual random forest classifiers. Confusion matrix and Matthews Correlation Coefficient (MCC) for each classifier.

The classification results were used to separate swing and stance phases. We defined the position of the tibia-tarsus joint in the last frame with ground contact as the posterior extreme position (PEP) and the first frame with ground contact after touch down as the anterior extreme position (AEP). All step parameters used in further analysis were calculated using these PEPs and AEPs. Legs were categorized as ‘inner’ and ‘outer’ legs depending on the sign of the radius of curvature of a given trajectory split (for details see section 2.4 Data Analysis). Left curves and corresponding step data were flipped and pooled together with right curves for all subsequent analysis.

### Step parameters and step type classification

Throughout this paper, we will refer to a step the sequence of beginning with an episode with “no ground contact” (e.g., swing) and a subsequent episode “with ground contact”. We describe either component by means of their length (in mm), duration (in s) and direction (in °), and implicitly assume that, on average, the parameters length and direction of the episode with “no ground contact” is opposite to those of the episode “with contact”. Generally, we focus on the episode with “no ground contact”, e.g., swing phases. Note that a leg movement with “no ground contact” could have different functions than a typical swing or return stroke. For instance, it could involve brief searching episodes. The “swing length” and “swing duration” of a step cycle were calculated as the distance and duration from PEP to AEP respectively (Fig. 3A). “Swing direction” was the direction of the vector between PEP and AEP (Fig. 3C). As a measure of how far and in which direction a leg was moved during a swing phase, we calculated the “swing angle” which we defined as the angular difference between the vectors from a thorax marker to the PEP and AEP of the respective leg (Fig. 3A). Compared to the swing direction, the swing angle does not have a cyclic nature as it can take on values only between ±90° during planar walking.

**Figure 3:**
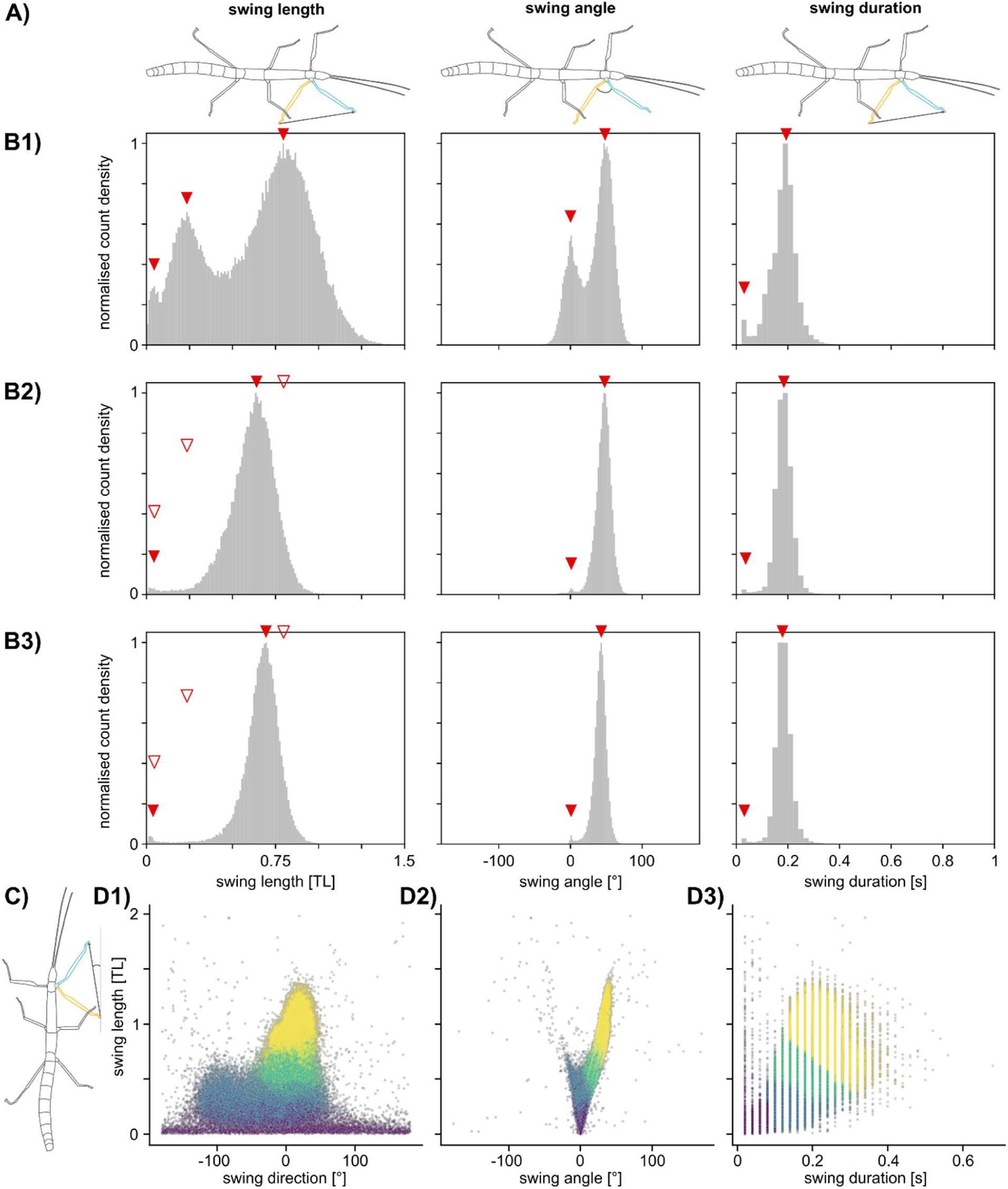
Data-driven K-Means clustering of step classes. **(A1-3)** Schematic illustrations of step parameters used for clustering. Blue leg is at AEP and orange leg at PEP. **(A1)** Swing length as a factor of thorax length (TL). **(A2)** Swing angle in degrees. **(A3)** Swing duration in seconds. **(B1-B3)** Normalised count density distributions of the three parameters (illustrated in A), as used for clustering. Red filled arrows point at local maxima of the respective distribution, red empty arrows point at local maxima of the front leg distribution. **(B1)** Front legs. **(B2)** Middle legs. **(B3)** Hind legs. **(C)** Swing direction. This parameter was not used in the K-Means algorithm but to evaluate whether step classes were sufficiently distinct and showed similar properties as described by Theunissen and Dürr (2013). The blue leg is at the AEP and the orange leg at the PEP. **(D1-3)** K-Means results. Cluster 1 (purple) is labelled as ‘short steps’, Cluster 2 (blue) as ‘middle steps’, and the union set of Clusters 3 (mint) and 4 (yellow) as ‘long steps’, yielding three step classes for subsequent analysis.

Swing length distributions of front legs were clearly not unimodal distributions, as observed previously by Theunissen and Dürr (2013) (Fig. 3B1), but this non-unimodality was less clear for middle or hind legs (Fig. 3B 2-3). Other than Theunissen and Dürr’s distinction of ‘short steps’ and ‘long steps’ on the grounds of the local minimum between the two modes of the swing length distribution, we opted for a data-driven method of step class differentiation. To this end, we used the K-Means clustering algorithm from scikit-learn to classify steps according to their swing length, swing angle and swing duration. Data from all legs was scaled using the *RobustScaler* function from scikit-learn. Other than in Theunissen and Dürr (2013), no probability distribution functions were fitted to the empirical distributions; neither was the classification done separately for each leg type. Instead, we trained a single classifier for step type classification irrespective of leg type. Finally, other than earlier step type classification, our method involved three step parameters (length, angle and duration) instead of only one (length). Since the mean swing length was smaller for middle and hind legs than for front legs, we needed to account for a potentially separate ‘long step’ cluster for these leg types, as opposed to that for front legs. For this we included the mean of the combined distribution of middle and hind legs as a separate prior. Accordingly, the algorithm was initialized with four sets of weights, which corresponded to the four peaks of overall swing parameter distributions shown in Fig. 3B 1-3.

The four resulting clusters differed strongly with regard to the three parameters (Fig. 3D 1-3). Cluster 1 (purple) had the shortest swing length and duration but varied strongly in swing direction: swing direction was distributed uniformly without an evident preferred direction. This was different from the other clusters, but similar to the properties of short steps as described by Theunissen and Dürr (2013). Therefore, we classified steps within this cluster as ‘short steps’. Clusters 3 (mint) and 4 (yellow) did not vary much besides in their mean swing length and swing angle. Moreover, both showed similar swing directions as the long steps reported by Theunissen and Dürr (2013). Since the absolute size of the swing angle is largely dependent on swing length, and the two different clusters were strongly associated with distinct leg types (cluster 3 was more likely for middle and hind legs, whereas cluster 4 more for front legs, see Suppl. Fig. 3), we decided that these two clusters were not sufficiently distinct to be considered separate step classes. Accordingly, clusters 3 and 4 were merged into a single step class termed ‘long steps’. The steps with a medium swing length (cluster 2, blue) had a considerably different swing direction range and swing angle sign compared to steps with longer swing lengths but were also different to the short step class. Therefore, we created a third step class which we, from now on, refer to as ‘middle steps’.

### Data analysis at the step level

All subsequent analysis was done for front legs only, but see Suppl. Mat. 4 for a comparison with middle and hind legs. After classifying all steps into three classes, we quantified their occurrence and potential changes in swing direction during straight walking as opposed to turning. To this end, we separated curved from straight trajectory segments according to their ROC.

To compare whether the three step types occurred at different frequencies for inner and outer legs (with regard to the curvature), and whether this changed as a function of curvature, we introduced an asymmetry measure as the dot product A=(S_inner_, M_inner_, L_inner_)·(S_outer_, M_outer_, L_outer_), where S, M and L are the percentages of short, middle and long steps, respectively. Percentages were calculated as means per animal for the two three-second episodes A and B (Fig. 1).

To address the question whether leg movements were adapted as a function of total turn angle or rather of maximum curvature we compared turning responses to 60° or 120° displacements in landmark position. For this, we calculated the change in swing direction as well as the change in its variance, using the circular summary statistics “mean swing direction” and “length of the resultant mean direction vector”, again as means per animal and for both episodes A, and B. To compare the latter, we calculated the mean differences of the circular summary statistics.

To compare the frequency of short steps during spontaneous and visually guided turns, and to relate this to short step frequency in earlier studies, we calculated the fraction of short steps for trajectory segment of a given range of ROC (see section 2.4). Again, this was done as means per animal per bin. For statistical comparison between trials with and without a landmark, i.e. visually guided as opposed to spontaneous turns, we used a Friedman test across all bins to test the null hypothesis that both conditions were equal. This was followed by pair-wise post-hoc Wilcoxon signed-rank tests for comparisons within bins.

## Results

### Path length, curvature and forward velocity differ among conditions

Animals readily and reliably approached the visual landmark, i.e. the black bar, if present (Fig. 1, Fig. 2, Fig. 4). The stepwise change in landmark position did not decrease the likelihood of an animal to approach the wall or landmark (Suppl. Tab. 1). However, for the larger displacement, the average likelihood to approach the target decreased by 20 percent points compared to the smaller displacement (Fig. 4B, Suppl. Tab. 1). In the all-white condition, animals approached the arena wall less often compared to trials in which a landmark was present (Fig. 4B, Suppl. Tab. 1). Without visual guidance, animals tended to walk around in circles (Fig. 2A, Fig. 4A). Trajectories in the all-white condition sometimes comprised one or multiple 360 degree turns (for example, see Fig. 1C). Overall, the probability of an animal to be *’on target*’ or *’at wall’* seemed to be unaffected by both the starting position of the animal in the arena and the direction of the landmark displacement (Fig. 4B, C, Suppl. Tab. 1). Therefore, we decided to pool both starting positions and displacement directions for the remaining analysis. Animals did not show a persistent orientation towards the original landmark position in trials with a change in landmark position, but re-oriented towards the new target instead (Fig. 4A, B). For all-white trials, we found no support for a preferred final position as in landmark trials.

**Figure 4:**
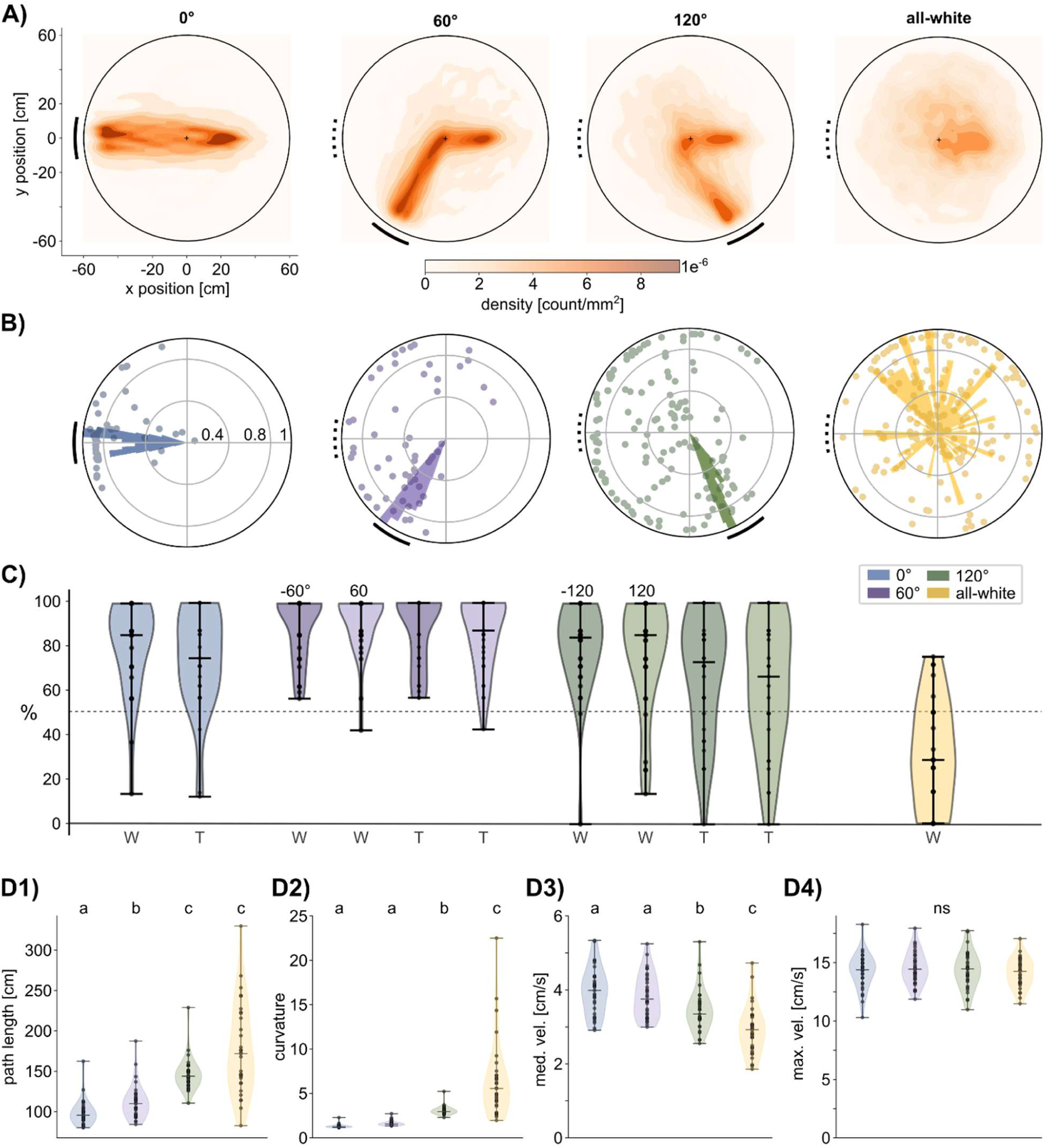
Final head position and likelihood of approach depends on presence of landmark. **(A)** Kernel density estimation plots across all trials. Solid black lines outside the circle show the width and position of the black bar presented. Dotted black lines show the first position of the black bar before it was displaced in 60° and 120° trials. In the all-white trial, the dotted line indicates the start orientation of the animal, since no black bar was present at any time. **(B)** Rose plots show the final head position of all trials that terminated at the arena wall (‘at wall’ trials). Colored circles show the end positions of trials in which animals were not ‘at wall’. **(C)** Violin plots show the likelihood of an animal to be ‘at wall’ (W) and ‘on target’ (T). Each data point represents one mean per animal (N = 31). ±60° and ±120° trials are shown as separate violin plots but were pooled for rose plots above. The detailed statistical results can be found in Suppl. Tab. 1. **(D1-4)** Violin plots show the mean across on-target trials of 30 individual animals of three different measures. Each data point represents one mean per animal. **(D1)** Path length. **(D2)** Curvature. **(D3)** Median forward velocity. **(D4)** Maximum forward velocity. Statistical results as well as mean and s.e.ms can be found in Suppl. Tab. 2.

Generally, the overall curvature of a trial was increased for all-white trials compared to trials with a landmark present (Fig. 4D2, Suppl. Tab. 2). The 60° condition did not differ from the 0° condition regarding median forward velocity, maximum forward velocity or overall curvature present (Fig. 4D, Suppl. Tab. 2). Contrary, the 120° and all-white conditions both showed an increased curvature and a decreased median forward velocity (Fig. 4D2, 3, Suppl. Tab. 2). The maximal forward velocity was not affected by a change in landmark position or the absence of a landmark present (Fig. 4D4, Suppl. Tab. 2).

### Displacement of landmark affects turning duration more than rotational velocity

To address the question how turning velocity and duration scale with the amplitude of a turn, we compared three measurements: the maximum rotational velocity and its timing, as well as the duration of the turn. Since forward velocities have been reported to decrease or increase during turning, depending on the type of turn (e.g. Yang et al. (2024)), we analyzed whether rotational and forward velocities followed a similar time course and whether forward velocity also scaled with the amplitude of a turn. Overall, we found that trials in which the landmark was moved by 120° appeared more variable in the approach of the new landmark position. This was particularly visible in the spatial distribution of episode B which covered the largest rotation throughout a trial (Fig. 2A, Fig. 5A2). We separated trials in which animals turned away from the target and approached it coming from the other side. These ‘away’ trials were more abundant in 120° trials (Fig. 5B). To further constrain the comparison to similar response types, we distinguished three categories according to where the maximal rotational velocity occurred: close to, intermediate (middle) and far from the midline of the arena. Again, the number of trials falling in to these three categories varied between 60° and 120° turns. While animals turned close to the midline in 90 percent of the 60° trials, they did so in only 46 percent of 120° trials. The time courses of rotational velocities varied considerably for the three different categories for 120° trials. Close turns showed a fast increase in rotational velocity followed by a short plateau at maximum rotational velocity and a fast but gradual decrease. In middle turns, animals rotated throughout the whole trial. In far turns, the onset of rotation was delayed by approximately three seconds (Fig. 5C, right column).

**Figure 5:**
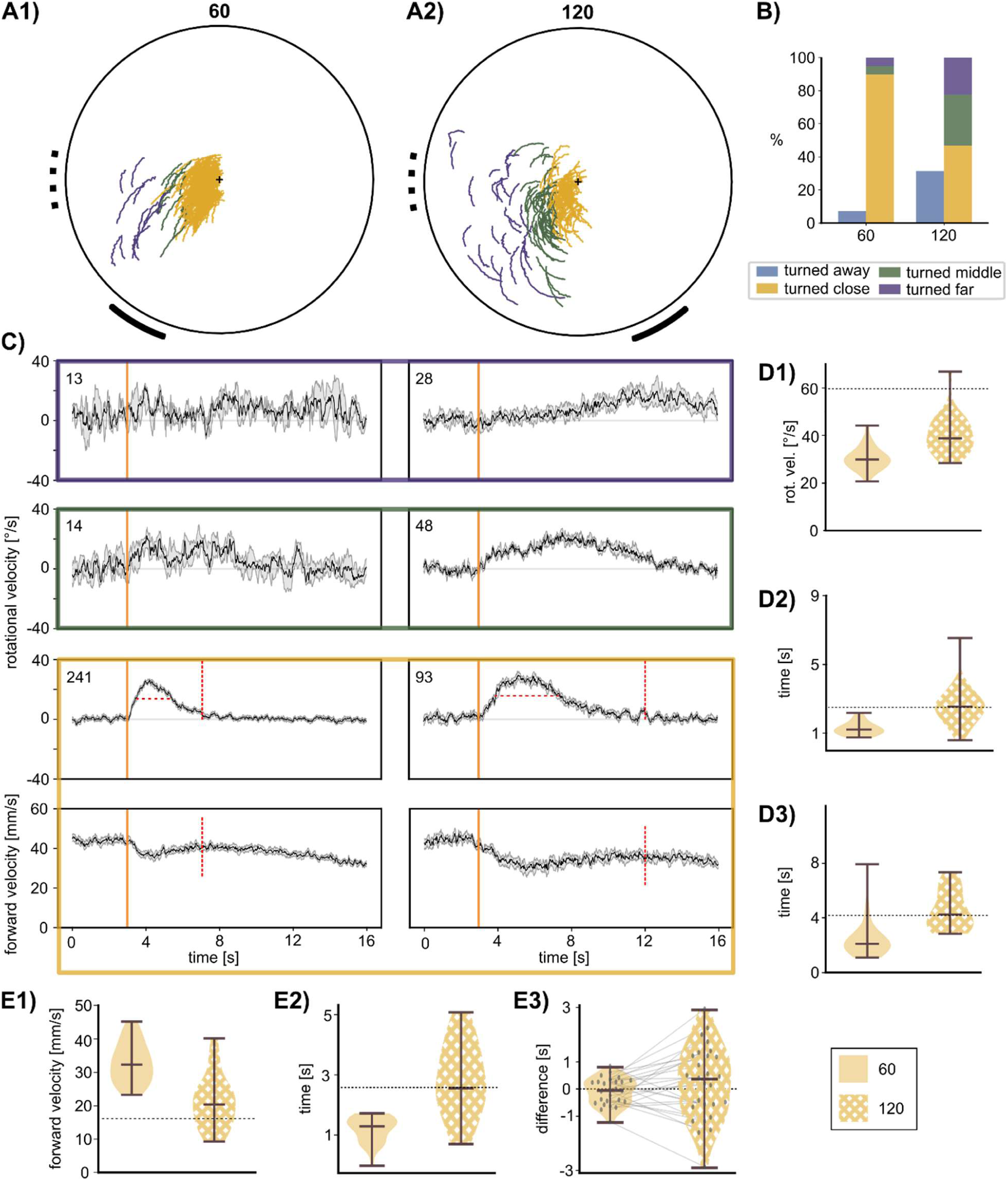
Maximum rotational velocity is larger and occurs later with a larger landmark displacement. **(A)** Top view of circular arena with xy-positions of the prothorax marker during episode B. All ‘central’ trials with turns towards the displaced landmark are shown. Solid black lines outside the circle show the width and position of the landmark presented. Dotted black lines show the initial position of the landmark before it was displaced. Plot only includes ‘on target’ trials. The black cross indicates the midpoint of the arena. Trajectories are color-coded according to the location of maximal turning. **(B)** Bar plots show the percentage of ‘central’ trials in which an animal turned away from the new landmark position (blue), turned close to the midpoint (yellow), turned in an intermediate range (green), or turned far away from the midpoint (purple). **(C)** Bootstrapped rotational velocity time courses. Black lines show the mean and grey lines show the 95 percent confidence intervals around the bootstrapped mean. Numbers on the upper left indicate the number of trials used for bootstrapping. Colored boxes around the plots correspond to the category of their turn location, as in (B). For ‘close towards’ trials (yellow) the second plot also shows the forward velocity time course. Horizontal red, dashed lines schematically show the FWHM. Vertical red, dashed lines show the time interval used for the calculation of minimum forward velocity. **(D1)** Violin plots show the maximum rotational velocity, **(D2)** the timing of maximum rotational velocity after the displacement of the landmark and **(D3)** the width at half maximum as a measure of rotation duration. To assess parameter scaling with turn angle, dotted lines indicate twice the median value of 60° turns. **(E1)** Violin plots show the minimum forward velocity, **(E2)** the timing of minimum forward velocity after the displacement of the landmark, and **(E3)** the temporal difference between the minimum forward velocity and maximum rotational velocity. Grey dotted line indicates zero-time difference. Grey dots show individual data points and grey solid lines connect data points of the same animals in both conditions. Values used for statistical comparison were means per animal. N = 29.

The quantitative comparison of 60° and 120° turns was confined to close turns. To distinguish between the alternative hypotheses that turns with twice the turn angle involved either two-fold rotational velocity (Fig. 5D1) or double the duration, we compared the mean parameters per animal against the theoretical predictions for peak rotational velocity (Fig. 5D1), time of peak velocity (Fig. 5D2), and the turn duration at more than half the peak velocity (full width at half max.; Fig. 5D3). On average, animals turned for nearly twice as long in 120° turns compared to 60° turns (p < 0.0001, mean ± s.e.m.: 60°: 2.44 ± 0.26 s, 120°: 4.57± 0.27 s, Fig. 5, D3), suggesting that animals covered a larger turn angle by increasing turn duration rather than speed. However, maximum rotational velocity was increased too for larger displacement magnitudes, albeit with mean velocities well below the predicted double velocity (p < 0.0001, mean ± s.e.m.: 60°: 30.35 ± 0.91 °/s, 120°: 40.26 ± 1.54 °/s Fig. 5D1). In addition, the maximum rotational velocity was reached later in 120° trials (p < 0.0001, mean ± s.e.m.: 60°: 4.28s ± 0.07s, 120°: 5.57s ± 0.21s, Fig. 5, D2). We conclude that our results support the hypothesis that larger turn angles are covered by proportionally longer turns (here: a factor of 1.87 compared to an expected 2.0), but that the scaling of turn parameters is non-linear. This non-linearity results in increased mean rotational velocities by a factor of 1.32.

For both turn angles, forward velocity decreased alongside the increase in rotational velocity (Fig. 5C). Here, minimum forward velocities also scaled non-linearly with turn angle as forward velocity was decreased more for larger turn angles by a factor of 1.54 (p < 0.0001, mean ± s.e.m.: 60: 32.98 mm/s ± 1.18 mm/s, 120: 21.35 mm/s ± 1.49 mm/s, Fig. 5E1). Both minimum forward and maximum rotational velocity share a similar timing, however they did not occur at the exact same time (see individual datapoints in Fig. 5E3). The difference of the timing of minimum forward velocity and maximum rotational velocity did not differ from zero for either displacement magnitude (p_60_ = 0.34, p_120_ = 0.51, mean ± s.e.m.: 60°: -0.11 s ± 0.09 s, 120°: 0.2 s ± 0.26 s, Fig. 5E3). However, as the distributions of difference in timing were nearly symmetrical around zero, albeit never being zero, the minimum forward velocity was either followed or preceded by the maximum rotational velocity. As before, 120° turns were more variable than 60° turns (Levene’s test: p < 0.0001, Fig. 5E3). Especially for 60° turns, the timing of these two events was less than a second apart for the majority of the animals, indicating a linkage between rotational and forward velocity albeit with two different relationships: Maximal rotational velocity leads minimum forward velocity or vice versa. These ‘strategies’ were not consistent for individual animals (Wilcoxon signed-rank test: p = 0.0005, indicated by zero crossings of grey lines in Fig. 5E3) revealing that animals changed strategies between different trials. In summary, we found that both, rotational and forward velocity scale with larger turn angles in a non-linear manner and that both parameters share a similar time course. However, there was no evidence for a strict coupling of forward and rotational velocity during turning.

### Presence of a visual landmark increases velocity and curvature

Our second objective was to test whether spontaneous turns differ from visually guided turns. Like guided turns, spontaneous turns varied in curvature. Accordingly, we addressed the question whether visual guidance in form of a landmark goes along with an increased forward velocity for similar curvatures. To this end, we divided each trial into segments of different curvature and compared forward velocities between episodes of equal curvature with or without visual guidance.

First of all, visual guidance in form of a black landmark increased the number of straight segments within a trial (Fig. 6). While 83.4 % of segments in all-white trials (Fig. 6B) were within a radius of curvature (ROC) of ± 600mm, only 58.1 % of guided trials (Fig. 6 A) fell into that range (for visualization of ROC, see Suppl. Fig. 2). As in previous results, forward velocity varied largely among animals (Fig. 6C, Fig. 3D3) and was linked to curvature (Fig. 5E1).

**Figure 6:**
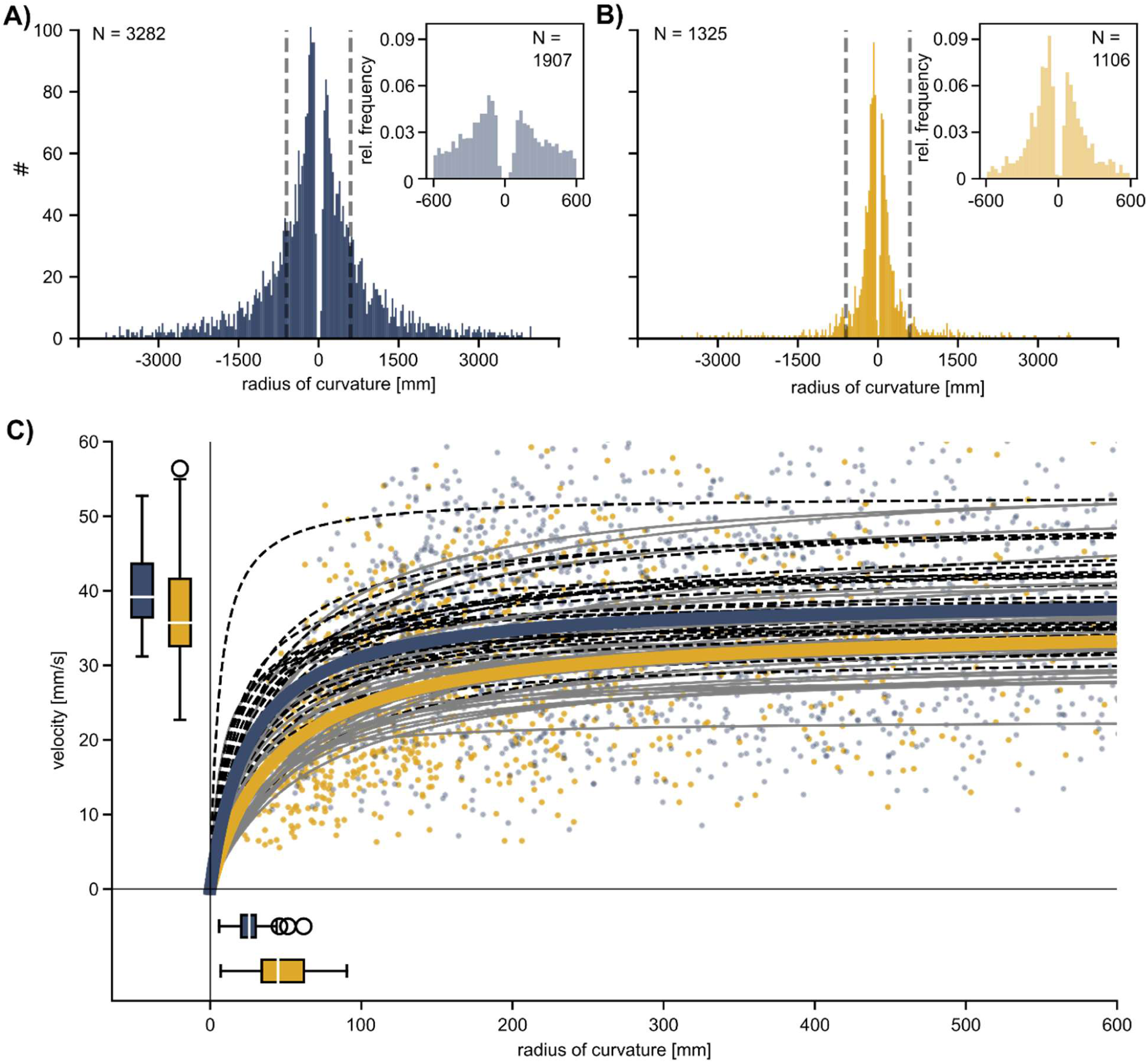
Guided turns are faster than spontaneous turns with the same radius of curvature. **(A, B)** Histograms show the number of trajectory segments with a given radius of curvature (ROC) for guided (A, blue) and spontaneous (B, yellow) turns. Inserts on the upper right are zoomed-in relative frequencies of the area marked by the vertical, dashed grey lines. 55 % of all segments were clockwise turns (right), the remaining 45% were counter-clockwise turns (left). Bin width = 30 mm; N = 30 animals. **(C)** Estimated saturation functions per animal are shown by grey solid lines (spontaneous condition) and black dashed lines (guided conditions). Thick blue and yellow lines show pooled saturation functions using the median maximum forward speed (V_max_) and the radius of curvature (ROC) at half V_max_ (K_M_) for guided and spontaneous conditions, respectively. Blue and yellow dots in the background show values of all trajectory segments clipped to ROC < 600 mm and V_max_ < 60 mm/s. Note that saturation curves were fit to non-clipped data ranges.

To quantify the curvature-velocity dependence we fitted separate saturation functions for each animal and statistically compared the maximum forward speed (V_max_) and the ROC at half V_max_ (K_M_). We found that both, K_M_ and V_max_, differed between the spontaneous and guided condition (p_VMax_ = 0.03, p_KM_ < 0.0001). Without visual guidance, the average maximum forward velocity was decreased (V_max_ guided: 40.27 ± 1.01, V_max_ spontaneous: 38.05 ± 1.43; mean ± s.e.m) and the ROC at half V_max_ was increased (K_M_ guided: 27.33 ± 2.25, K_M_ spontaneous mean ± s.e.m.: 47.99 ± 3.86; mean ± s.e.m), indicating that curves of similar curvature were walked at a higher forward velocity if a landmark was present (Fig. 6B).

### Stick insects use three classes of steps during unrestrained locomotion

To address our questions at the level of single steps, we first had to detect and classify all steps. Although this was done for all six legs, our hypotheses focused on front legs in particular, such that we will report results on front legs only. All material concerning middle and hind legs can be found in the Supplementary Material (e.g., Suppl. Fig. 4).

Generally, we found strong variation of step length during unrestrained turning on a planar surface. Associated step length distributions were not unimodal, suggesting the presence of distinct step classes. However, step length distributions differed from those found in studies that used “straight walkway” setups (Theunissen and Dürr, 2013; Theunissen et al., 2015). Instead of a bimodal distribution with short and long steps, we found a trimodal distribution of swing length (Fig. 3 B1). To check whether the medium swing length mode was related to turning rather than to straight walking, we first separated steps from nearly ‘straight’ segments, i.e., with a ROC larger than 600 mm, from ‘curved’ segments with a smaller ROC. Since steps of medium length occurred in straight and curved segments alike, though with different frequencies (Fig. 7A, C), we concluded that a binary distinction between long and short steps was inappropriate for our data. Following data-driven categorization based on the three step parameters length, duration and angle (Section 2.6, Fig. 3), we found that steps could be divided into three step classes that corresponded to the three modes of the step length distribution: short, middle and long steps.

**Figure 7:**
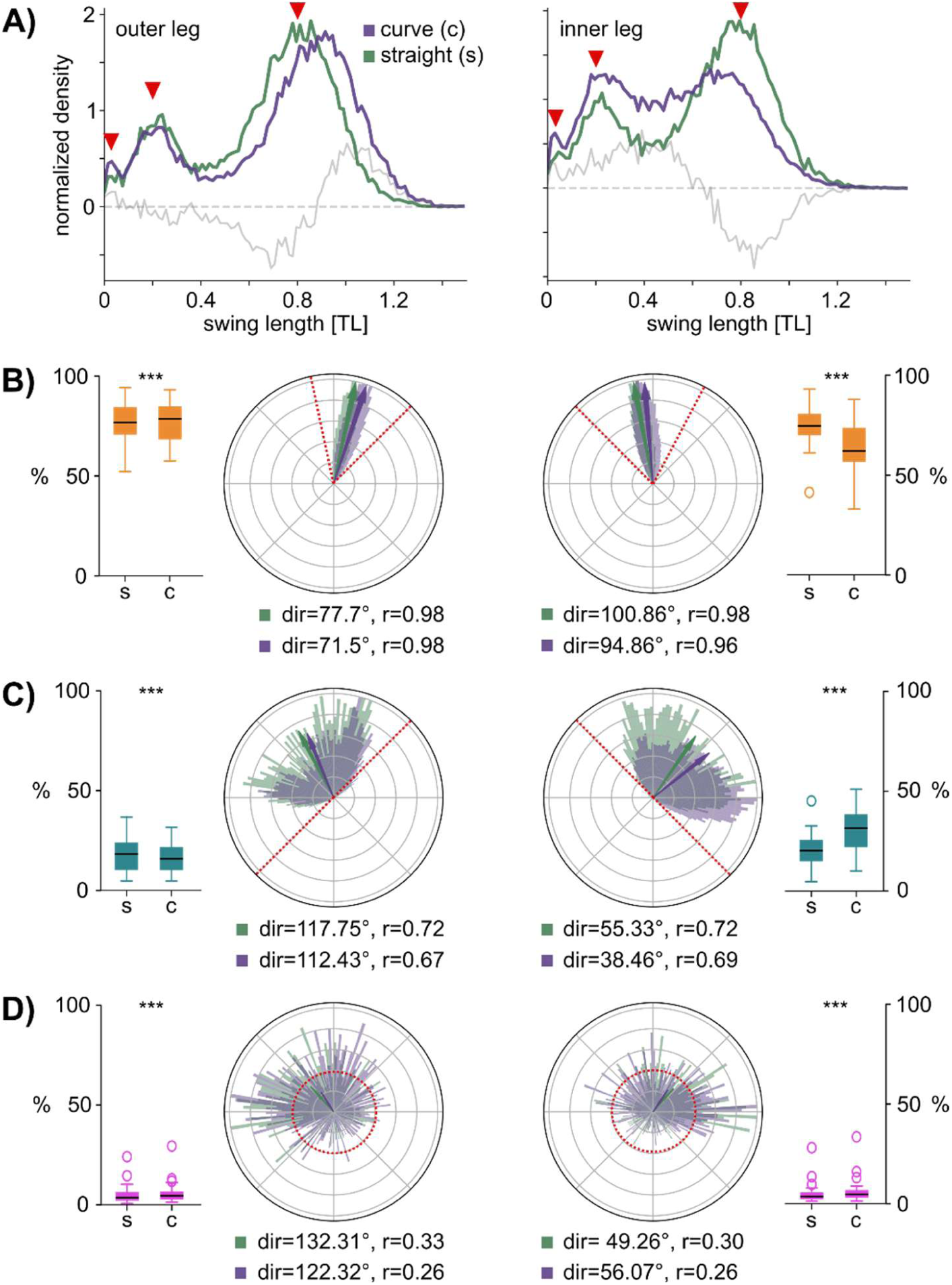
Stick insects use three classes of steps during unrestrained planar walking. (**A**) Density plots show the distribution of swing lengths of front legs (left: outer leg, right: inner leg) in curved (purple, ROC <= 600, N=32, n_steps_inside_= 25202, n_steps_outside_= 21896) and straight segments (turquoise, ROC > 600 mm, N=32, n_steps_inside_ = 16375, n_steps_outside_ = 15951). The grey solid line shows the difference between the curved and straight conditions. The grey dashed line marks zero density or difference. N = 32 (**B-D**) Circular histograms show the swing direction of front leg steps (left column: outer leg, right column: inner leg) in curved (purple) and straight (green) segments. The mean direction (dir) and the concentration around the mean direction (r) are given below. (**B**) Long steps. (**C**) Middle steps. (**D**) Short steps. Boxplots show the percentages of a step type for straight (S) and curved (C) segments; N = 31 means per animal. For statistical results see Suppl. Tab. 3.

When comparing the properties of these three step classes, swing directions of short and long steps were similar to findings of previous studies: While long steps were directed towards the front with a slight medial slant (Fig. 7B), short steps had little to no directional bias (Fig. 7D). Other than that, middle steps were directed laterally and, similar to long steps, slightly medially. This was the case in both, inner and outer legs. However, unlike short steps, they never pointed into the medio-caudal half of the unit circle. The variance in swing directions was larger for middle steps than for long steps, but smaller compared to short steps (as reflected by mean vector length r in Fig. 7B-D). During curve walking, the mean swing direction of long steps was shifted towards the direction of the curve for both, inner and outer legs. In middle steps, this shift in mean direction was stronger for the inner leg. Here, also the frequency of middle steps increased during curve walking, while the percentage of long steps decreased accordingly (see boxplots in Fig. 7B-D). The frequency of all three step classes changed during curve walking (Suppl. Tab. 3).

Additionally, we found that left-right asymmetry of long and middle step frequencies was prominent during curve walking where inner legs take fewer long steps and more middle steps compared to outer legs (long steps: -1.61, middle steps: 1.84, Cohen’s δ). We found a similar relationship for near ‘straight’ walks, although with low effect sizes (long steps: -0.27, middle steps: 0.31, Cohen’s δ) as this effect is likely linked to the fact that ROCs of 600mm still being shallow curves. Taken together, we found that step type frequencies for all legs change from straight to curve walking as well as an asymmetry in step type frequencies between inner and outer legs during curve walking.

### Swing directions are altered more strongly for larger turn angles

As the previous result already indicated that swing direction as well as step type frequencies changed during curve walking, we wanted to find out how the scaling of turning responses is carried out at the level of individual steps. Accordingly, we calculated the change in swing direction as well as the change in variance of swing directions from the ‘straight walking’ episode A to the ‘curve walking’ episode B, as distinguished in Figs. 1, 2. To relate this to the body kinematics of Fig. 5, we only used ‘central’ trials which were categorized as ‘close’ turns ‘towards’ the landmark.

For both types of trials, turn angles were the same during ‘straight’ episodes A, and significantly increased during ‘curved’ episode B (Fig. 8A). Throughout the three seconds of ‘curved’ episode B, animals covered an approximately 50 % larger turn angle in 120° trials compared to 60° trials (Fig. 8A). The different turn angle between straight and curved episodes was accompanied by a corresponding change in left-right asymmetry of step type frequencies, as reflected by the smaller dot product of the frequency vectors for inner and outer legs (Fig. 8B; see section 2.7). Generally, left-right asymmetry increased during turning. It was significantly larger for 120° trials than for 60° trials (Fig. 8B).

**Figure 8:**
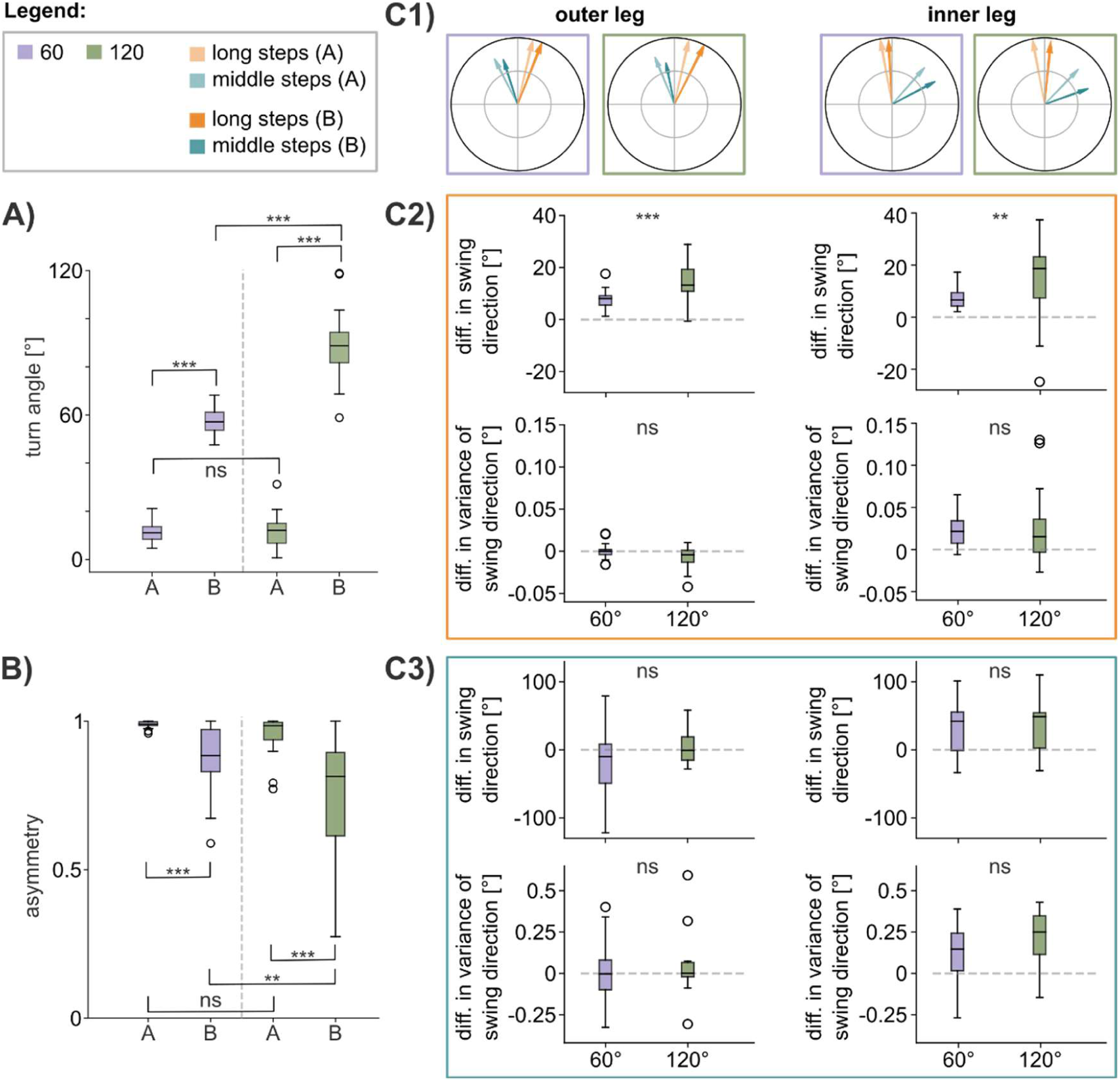
Swing direction changes are larger during prolonged rotation. (**A**) The turn angle covered during 3 s episodes A and B differs between 60° (purple) and 120° (green) trials. (**B**) Asymmetry between step class frequencies of inner and outer front legs during straight episode A (left) and curved episode B (right) differs between60° (purple) and 120° (green) trials. (**C1**) Median swing direction across all animals of two step classes (orange: long steps, turquoise: middle steps) for both front legs (left: outer leg, right: inner leg) during 60° (purple) and 120° trials (green). Arrows of lower opacity belong to the straight episode A, those of higher opacity to curved episode B. Inner grey circle shows r = 0.5. Outer black circle shows r = 1. N=29. (**C2**) Upper row of plots show the difference in swing directions of long steps between episodes A and B as means per animal for outer leg (left) and inner leg (right). Each plot compares 60° (purple) against 120° trials (green). Lower row of plots shows the difference in variance of swing directions of long steps between episodes A and B, as means per animal (left: outer leg, right: inner leg). Each plot compares 60° (purple) against 120° trials (green). (**C3**) Same as C2 but for middle steps. For statistical results and all mean and s.e.m. values, see Suppl. Tab. 4-6.

During turning, swing direction changed significantly more in 120° trials compared to 60° trials for both inner (Fig. 8C2; 60°: 7.07 ± 0.63°, 120°: 14.7 ± 2.62°; mean ± s.e.m.) and outer legs (60°: 7.91 ± 0.61°, 120°: 14.38 ± 1.24°; mean ± s.e.m.). Swing directions of the inner leg were directed further towards the inside of the curve, with a more than two-fold larger angle in 120° trials (factor 2.08). In outer legs, the increase was also significant, though slightly smaller (factor 1.82; Fig. 8C2). The turning-related change in variance of swing direction among long steps was larger in inner legs compared to outer legs for 60° trials (inner: 0.0243 ±0.033, outer: -0.0007 ± 0.0014; mean ± s.e.m.) and for 120° trials (inner: 0.0226 ± 0.0071, outer: -0.0062 ± 0.0022; mean ± s.e.m.; Fig. 8C2) but similar between the two conditions.

Other than long steps, the turning-related change in swing direction of middle steps did neither differ between inner and outer legs, nor between 60° and 120° turns (Fig. 8C3). Although the variance of swing direction changes was very large for both turn angles, it did not differ significantly (Fig. 8C3).

Overall, we found that during turning long steps of inner and outer legs were both rotated towards the direction of the turn. This effect was nearly twofold larger for 120° turns compared to 60° turns. At the same time, swing direction of middle steps did not change (Fig. 8C), despite a turning-related change in step type asymmetry between inner and outer legs (Fig. 8B).

### Front legs execute more short steps during tight turns without visual guidance

Finally, to test whether short steps might be used as intermitted ‘searching steps’ in a walking scenario without visual guidance, we compared the relative frequency of short steps between trials with visually guided turns and trials with spontaneous turns, i.e. between trials with and without a visual landmark, respectively. To account for different radius of curvatures, we used four bins of ROC from the segment analysis in Figure 6.

Overall, we found a small but statistically significant increase in the relative frequency of short steps in all-white trials compared to visual landmark trials (Fig. 9, Friedman test: p<0.0001). Post-hoc paired tests for different ranges of ROC revealed that this significant difference was mainly carried by differences between tight turns. In strongly curved segments (ROC < 200 mm), short steps were about 1.5 percent points more frequent in all-white trials than in landmark trials (mean ± s.e.m.: steps_stim_= 4.6% ± 0.57%, short steps_white_ = 6.08% ± 0.73%, p = 0.0024). For shallower curves and straight segments, the number of short steps was not different for the two conditions (after Bonferroni correction).

**Figure 9:**
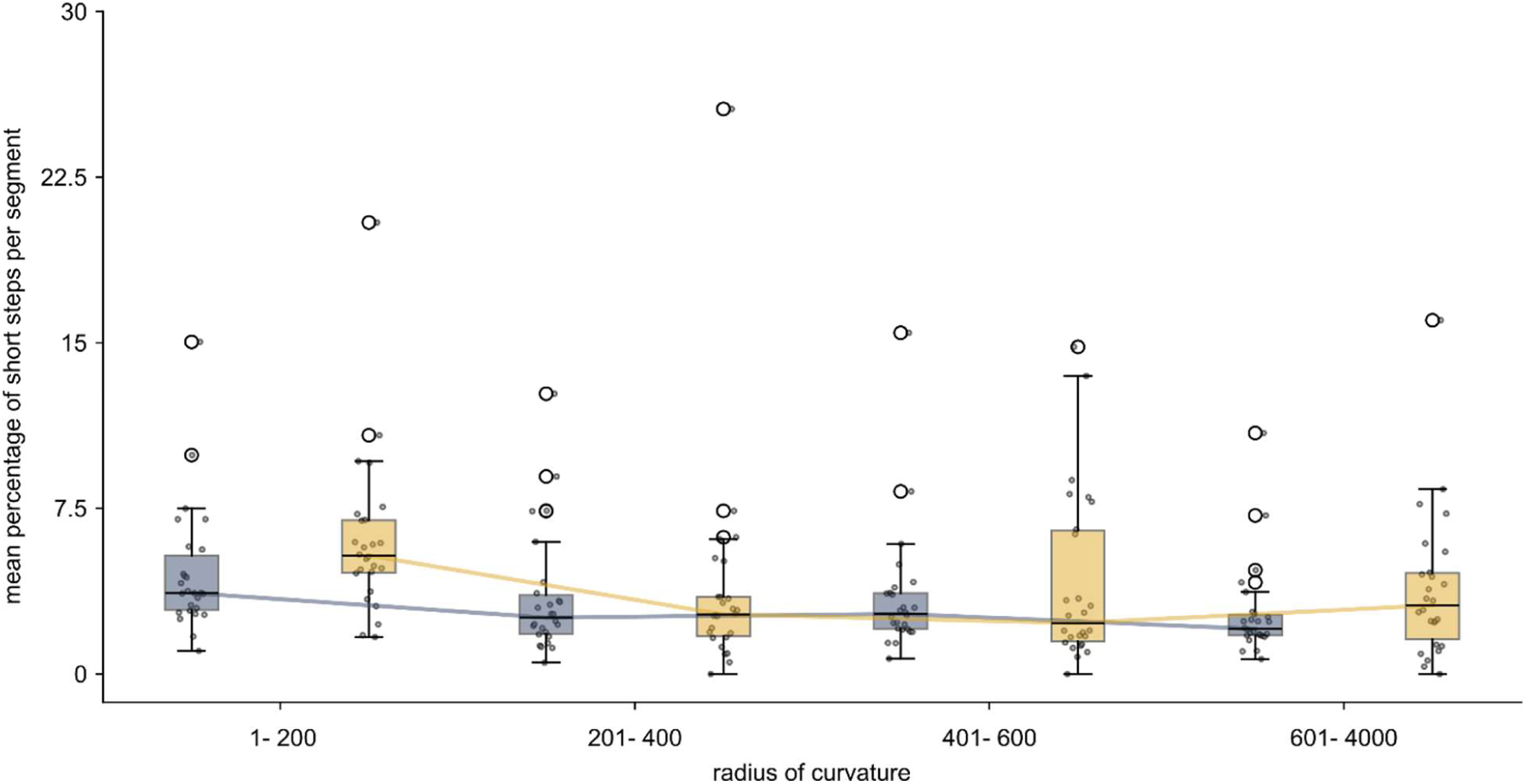
Short steps are more frequent in spontaneous tight turns. Boxplots show the percentage of short steps for path segments of four different radius of curvature (ROC) ranges in spontaneous (yellow) and visually guided (blue) curves. Segments were grouped according to their ROC as means per animal per condition; N = 26. For statistical results and all mean and s.e.m. values see Suppl. Tab. 7.

## Discussion

Flexible walking behaviour allows animals to change their heading according to context and across a range of curvatures. Here, we investigated contextual effects in the form of visual guidance, and magnitude-related effects in the form of scaled turning responses. For both of these aspects, we linked the behavioural level of body trajectories to the causal level of step parameters (with a focus on front legs). To induce different turn magnitudes, we modified a landmark displacement paradigm introduced by Rosano and Webb (2007), constraining both onset time and desired turn angle of unrestrained walking stick insects. To our knowledge, this is the first study to address the scaling of turning responses towards stationary targets in unrestrained walking.

### Turning responses scale non-linearly with turn angle

For turning responses that started at a similar location (‘close’ condition), we found that turning duration and maximum turning velocity scaled with the displacement of a landmark in a non-linear manner, as turning duration was affected to a larger extent (1.87×) than maximum turning velocity (1.32×) (Fig. 5). Similar to findings in spiders (Land (1972), the rotational velocity was not constant between or within turns, indicating that the turn towards a stationary target is not simply scaled according to an error signal (e.g. target eccentricity).

Since the scaling factor for turning duration was larger than that for rotational velocity (Fig. 5), one might expect rather small changes in leg movement parameters because turning duration should correlate with the number of steps needed to cover the larger turn angle. However, we found that changes in swing direction of both front legs were up to twice as large (Fig. 8C), suggesting that even small increases in rotational velocity arise from strongly altered leg movements. In addition, as the left-right asymmetry of step types changes significantly for larger turn angles, the mix of step parameters was altered differently for inner and outer legs, depending on turn angle. This is consistent with findings by Yang et al. (2024), who observed different combinations of ‘leg gestures’ for turn types of different curvature. Indeed, these authors could relate narrow ‘pivot’ *versus* shallow ‘swerve’ turns in spontaneously turning *Drosophila* to activity modulation of identified descending interneurons. Though both turn types were identified through transient increases of rotational velocity, the associated steering maneuvers involved either an increase (for shallow turns) or decrease (for narrow turns) in forward velocity (Yang et al. (2024)). For *C. morosus*, Dürr and Ebeling (2005) found no change in forward velocity during visual-motion-induced turns in tethered walking, but a small but significant increase in sideward velocity. In contrast to both of these studies, we found no evidence for an increase in forward velocity but rather a scaled reduction of forward velocity, depending on overall turn angle. It is possible that these differences correspond to different walking conditions (free vs. tethered) and/or paradigms (spontaneous, motion-induced or target-induced, see below). Since differences in the timing of minimum forward and maximum rotational velocity did not occur in a fixed sequence there is no evidence for a strict coupling of forward and rotational velocity during turning. Rather, different animals reached their maximum rotational velocity either before or after reducing their forward velocity to a minimum (Fig. 5E3). Thus, different individuals show different strategies and tuning of turning parameters to walk the same curve. However, since individuals also changed strategies between 60° and 120° turns, idiosyncrasies as observed for straight walking in *Drosophila* (Godesberg et al. (2024)) are unlikely to be the driver of these different strategies. Rather, these findings suggest a motor flexibility during turning. Taken together, the number of parameters that change during target-induced turning and their different scaling with turn magnitude support the idea that locomotion models should combine more than one mechanism to govern turning. Moreover, these mechanisms need to be adaptive to account for continuous scaling of curvature at different walking speeds, while retaining flexibility for multiple possible combinations of parameter scaling to change in heading. Our study sets the stage for a detailed sequel analysis on changes in inter-leg-coordination (Dürr (2005), Rosano and Webb (2007); DeAngelis et al., (2019)) for turns of different radii as well as on the flexibility of transition from straight to curve walking.

### Distinct step classes during turning

Adaptive walking has been shown to involve proprioceptive feedback in addition to changes in centrally generated rhythmic motor patterns (for reviews, see Ritzmann and Büschges 2007, Büschges et al., 2008), in correspondence with descending input from brain neuropils (Cruz and Chiappe, 2023). In turning stick insects, the strength in inter-leg coupling has been shown to change differently for distinct leg pairs, with a left-right asymmetry as well as a pronounced effect on front leg coupling (Dürr 2005). A near-complete disruption of a six-legged gait occurs in tight on-the-spot turns, e.g. when turning on a twig (Cruse et al., 2009b). In account of possible gait disruption, and the special role of front legs, our present paper focuses on front legs only (but see Suppl. Figs S3 and S4). This is further justified by the occurrence of distinct step types (Theunissen and Dürr, 2013; Theunissen et al., 2015; Bläsing and Cruse, 2004b) the identification of which requires looked at the stochastics of step parameter distributions. While neglecting temporal coordination among legs, this approach has the benefits to include aspects of spatial coordination as well as being applicable to walking sequences where no distinct gait is identifiable.

Although step parameters of both swing and stance phase change with corresponding magnitudes during turning (Dürr and Ebeling, 2005), a recent study shows that turns toward static visual landmarks tend to be initiated by a change in swing direction (Bigge et al., 2026). Accordingly, our step parameter analysis focused on swing phase parameters. Contrary to the studies by Theunissen et al., we found a trimodal step length distribution, comprising a third, previously undetected step class (Fig. 7). If we had applied the same step length criterion to separate steps into two classes as done by Theunissen and Dürr (2013), we would have obtained long steps and a joint grouping of middle and short steps. Thus, it is likely that middle and short steps were pooled together in previous publications. The short step class occurs less frequently in our analysis compared to Theunissen and Dürr (2013). We attribute this to the mentioned pooling in previous studies, but also to effects related to structural properties of the experimental setup. Paradigm-related effects on step parameters have been discussed for free walking as opposed to tethered walking on a treadwheel or a slippery plate (Cruse and Bartling, 1995) as well as for walkways of different slope (Grabowska et al., 2012). As Theunissen and Dürr (2013) worked with a narrow walkway, foot positions were generally close to the edges of the walkway, which could have increased the frequency of short tactile probing movements and/or correction steps in order to maintain a stability. In addition, middle steps, which were more frequent in our data, could have been less frequent on the narrow walkway, as these steps were often directed laterally and could be related to balancing, sideways translation and turning. Indeed, the turning-related increase in step type asymmetry (Fig. 7) and the pronounced left-right asymmetry of middle step direction of inner and outer legs during guided turns (Fig. 8C) suggests that middle steps are an important means of the animal during turning. Similarly, the pronounced mediad steps of front legs observed in visual-motion-induced turning (Dürr and Ebeling, 2005; Gruhn et al., 2009) correspond best to the step class which we, for now, have called middle steps.

That being said, the bimodal distribution of middle leg step direction (Fig. 7C) and the similarity of one of these modes with the prevalent long step direction (Fig. 7B) may indicate that our step type classification is not ideal yet. Future work may need to include the complete limb kinematics to characterize middle steps in more detail. Similarly, a sequel study shall focus on the occurrence of particular step types in temporal sequence and gait analysis, complementing our present understanding of middle step function.

### Visually guided *versus* spontaneous turns

Spontaneous turning in unrestrained terrestrial locomotion has been repeatedly studied in open-field settings without visual landmarks and little or no visual-motion cues (Strauss and Heisenberg, 1990; Jindrich and Full, 1999; Yang et al., 2024). Here, we used an open-field arena with a white wall (condition ‘all-white’) as a reference condition to address the questions (i) whether stick insects constrain their motor flexibility in a context-dependent manner, as indicated by differences compared to visually guided turns, and (ii) whether they compensate for the lack of visual guidance by increasing tactile sampling, as indicated by the frequency of short steps. Spontaneous turns with a similar ROC were, on average, walked at lower speeds compared to visually guided turns (Fig. 6). At present, we cannot distinguish whether this is due to distinct turning strategies related to different initiation of turning, or rather due to a difference on arousal or motivation related to the absence/presence of external stimuli. While the increased frequency of short steps for spontaneous turns with high curvature (Fig. 9) speaks for the former, the overall lower forward velocity (Fig. 6) speaks for the latter. Future work may need to test whether and how turning-related changes in parameters of long and middle steps depend on forward velocity in the same way in visually guided and spontaneous turns. As all-white trials were, on average, walked at lower speed, it is possible that the primary effect of lacking visual guidance is the increase in curvature, in turn leading to a reduction in forward velocity. Our finding that short steps were more frequent in narrow curves only (Fig. 9) supports the interpretation that short steps serve as correction steps (Theunissen and Dürr (2013)), as static stability may be compromised increasingly more at increasingly tighter turns.

## Conclusion

Taken together, our results show that stick insects alter their turning behavior depending on the visual context, as revealed by reduced overall speed and increased frequency of short steps. When turning towards static visual targets, turning responses scaled non-linearly, where scaling was dominated by an increase in turning duration rather than maximum rotational velocity. Increased rotational velocity corresponded with reduced forward velocity. These changes were caused by larger shifts in step direction, as well as an increased asymmetry in step types between inner and outer legs, with increased frequency of middle steps on the inner side of the curve. In summary, our results suggest a mix of distinct turning strategies with both the scaling of step parameters and the likelihood of particular step class depending on overall turn angle.

## References

Arent, I., Schmidt, F. P., Botsch, M. and Dürr, V. (2021). Marker-less motion capture of insect locomotion with deep neural networks pre-trained on synthetic videos. Front. Behav. Neurosci. 15, 637806.

Berg, E. M., Büschges, A. and Schmidt, J. (2013). Single perturbations cause sustained changes in searching behavior in stick insects. J. Exp. Biol. 216, 1064–1074.

Bigge, R., Schwermann, N. and Dürr, V. (2026). Changing 3D heading direction in terrestrial locomotion: Effectiveness of visual landmarks and the onset of turning in stick insects. J. Comp. Physiol. A, 10.21203/rs.3.rs-9243999/v1

Bläsing, B. and Cruse, H. (2004a). Mechanisms of stick insect locomotion in a gap crossing paradigm. J. Comp. Physiol. A 190, 173–183.

Bläsing, B. and Cruse, H. (2004b). Stick insect locomotion in a complex environment: climbing over large gaps. J. Exp. Biol. 207, 1273–1286.

Büschges, A., Akay, T., Gabriel, J. P. and Schmidt, J. (2008). Organizing network action for locomotion: Insights from studying insect walking. Brain Res. Rev. 57, 162–171.

Büschges, A. and Ache, J. M. (2025). Motor control on the move: from insights in insects to general mechanisms. Physiol. Rev. 105, 975–1031.

Cruse, H. and Bartling, C. (1995). Movement of joint angles in the legs of a walking insect, *Carausius morosus*. J. Insect Physiol. 41, 761–771.

Cruse, H., Dürr, V., Schilling, M. and Schmitz, J. (2009a). Principles of insect locomotion. In Spatial Temporal Patterns for Action Oriented Perception in Roving Robots (ed. P. Arena and L. Patanè), pp. 43–96. Berlin: Springer.

Cruse, H., Ehmanns, I., Stübner, S. and Schmitz, J. (2009b). Tight turns in stick insects. J. Comp. Physiol. 195, 299–309.

Cruz, T. L. and Chiappe, M. E. (2023). Multilevel visuomotor control of locomotion in *Drosophila*. Curr. Opin. Neurobiol. 82, 102774.

DeAngelis, B. D., Zavatone-Veth, J. A. and Clark, D. A. (2019). The manifold structure of limb coordination in walking Drosophila. eLife 8, e46409.

Dürr, V. (2001). Stereotypic leg searching-movements in the stick insect: Kinematic analysis, behavioural context and simulation. J. Exp. Biol. 204, 1589–1604.

Dürr, V. (2005). Context-dependent changes in strength and efficacy of leg coordination mechanisms. J. Exp. Biol. 208, 2253–2267.

Dürr, V. and Ebeling, W. (2005). The behavioural transition from straight to curve walking: kinetics of leg movement parameters and the initiation of turning. J. Exp. Biol. 208, 2237–2252.

Dürr, V. and Mesanovic, A. (2023). Behavioural function and development of body-to-limb proportions and active movement ranges in three stick insect species. J. Comp. Physiol. A, 265–284.

Franklin, R., Bell, W. J. and Jander, R. (1981). Rotational locomotion by the cockroach *Blattella germanica*. J. Insect Physiol. 27, 249–255.

Frantsevich, L. I. and Cruse, H. (2005). Leg coordination during turning on an extremely narrow substrate in a bug, *Mesocerus marginatus* (Heteroptera, Coreidae). Journal of Insect Physiology 51, 1092–1104.

Fujiwara, T., Brotas, M. and Chiappe, M. E. (2022). Walking strides direct rapid and flexible recruitment of visual circuits for course control in *Drosophila*. Neuron 110, 2124–2138.e8.

Godesberg, V., Bockemühl, T. and Büschges, A. (2024). Natural variability and individuality of walking behavior in Drosophila. J. Exp. Biol. 10.1242/jeb.247878.

Grabowska, M., Godlewska, E., Schmidt, J. and Daun-Gruhn, S. (2012). Quadrupedal gaits in hexapod animals - inter-leg coordination in free-walking adult stick insects. J. Exp. Biol. 215, 4255–4266.

Graham, D. (1985). Pattern and control of walking in insects. Adv. Insect Physiol. 18, 31–140.

Gruhn, M., Zehl, L. and Büschges, A. (2009). Straight walking and turning on a slippery surface. J. Exp. Biol. 212, 194–209.

Jander, J. P. (1985). Mechanical stability in stick insects when walking straight and around curves. In Insect Locomotion (ed. M. Gewecke and G. Wendler), pp. 33–42. Hamburg, Berlin: Paul Parey.

Jindrich, D. L. and Full, R. J. (1999). Many-legged maneuverability: dynamics of turning in hexapods. J. Exp. Biol. 202, 1603–1623.

Kalmus, H. (1937). Photohorotaxis, eine neue Reaktionsart, gefunden an den Eilarven von *Dixippus morosus*. Z. vergl. Physiol. 24, 644–655.

Land, M. F. (1972). Stepping movements made by jumping spiders during turns mediated by the lateral eyes. J. Exp. Biol. 57, 15–40.

Mathis, A., Mamidanna, P., Cury, K. M., Abe, T., Murthy, V. N., Mathis, M. W. and Bethge, M. (2018). DeepLabCut: markerless pose estimation of user-defined body parts with deep learning. Nat. Neurosci. 21, 1281–1289.

Meschenmoser, M. and Dürr, V. (2025). Contrast and luminance dependence of target choice and visual orientation in walking stick insects. Scientific Reports 15, 12226.

Nath, T., Mathis, A., Chen, A. C., Patel, A., Bethge, M. and Mathis, M. W. (2019). Using DeepLabCut for 3D markerless pose estimation across species and behaviors. Nature protocols 14, 2152–2176.

Niemeier, M., Jeschke, M. and Dürr, V. (2021). Effect of Thoracic Connective Lesion on Inter-Leg Coordination in Freely Walking Stick Insects. Frontiers in bioengineering and biotechnology 9, 628998.

Nye, S. W. and Ritzmann, R. E. (1992). Motion analysis with leg joints associated with escape turns of the cockroach, *Periplaneta americana*. J. Comp. Physiol. 171, 183–194.

Pedregosa, F., Varoquaux, G., Gramfort, A., Michel, V., Thirion, B., Grisel, O., Blondel, M., Müller, A., Nothman, J., Louppe, G. et al. (2011). Scikit-learn: Machine Learning in Python. Journal of Machine Learning Research (.

Rayshubskiy, A., Holtz, S. L., Bates, A. S., Vanderbeck, Q. X., Serratosa Capdevilla, L., Rockwell, V. and Wilson, R. I. (2025). Neural circuit mechanisms for steering control in walking *Drosophila*. eLife. 10.7554/eLife.102230.3.

Ridgel, A. L., Alexander, B. E. and Ritzmann, R. E. (2007). Descending control of turning behavior in the cockroach, *Blaberus discoidalis*. J. Comp. Physiol. A 193, 385–402.

Ritzmann, R. E. and Büschges, A. (2007). Adaptive motor behavior in insects. Curr. Opin. Neurobiol. 17, 629–636.

Rosano, H. and Webb, B. (2007). A dynamic model of thoracic differentiation for the control of turning in the stick insect. Biol. Cybern. 97, 229–246.

Schütz, C. and Dürr, V. (2011). Active tactile exploration for adaptive locomotion in the stick insect. Phil. Trans. R. Soc. Lond. B 366, 2996–3005.

Sinéty, R. de (1901). Recherches sur la biologie et l’anatomie des Phasmes. La Cellule XIX, 118–278.

Strauss, R. and Heisenberg, M. (1990). Coordination of legs during straight walking and turning in *Drosophila melanogaster*. J. Comp. Physiol. A 167, 403–412.

The MathWorks Inc. (2022). MATLAB version: 9.13.0 (R2022b). Natick, Massachusetts, United States: The MathWorks Inc.

Theunissen, L. M. and Dürr, V. (2013). Insects use two distinct classes of steps during unrestrained locomotion. PLOS one 8, e85321.

Theunissen, L. M., Bekemeier, H. H. and Dürr, V. (2015). Comparative whole-body kinematics of closely related insect species with different body morphology. J. Exp. Biol. 218, 340–352.

Truong, C., Oudre, L. and Vayatis, N. (2020). Selective review of offline change point detection methods. Signal Processing 167, 107299.

Van Rossum, Guido and Drake, Fred L. (2009). Python 3 Reference Manual. Scotts Valley, CA: CreateSpace.

Virtanen, P., Gommers, R., Oliphant, T. E., Haberland, M., Reddy, T., Cournapeau, D., Burovski, E., Peterson, P., Weckesser, W., Bright, J. et al. (2020). SciPy 1.0: fundamental algorithms for scientific computing in Python. Nature Methods 17, 261–272.

Wilson, D. M. (1966). Insect walking. Ann. Rev. Entomol. 11, 103–122.

Yang, H. H., Brezovec, B. E., Serratosa Capdevila, L., Vanderbeck, Q. X., Adachi, A., Mann, R. S. and Wilson, R. I. (2024). Fine-grained descending control of steering in walking *Drosophila*. Cell 187, 6290–6308.e27.

Zeng, Y., Chang, S. W., Williams, J. Y., Nguyen, L. Y.-N., Tang, J., Naing, G. and Dudley, R. (2020). Canopy parkour: movement ecology of post-hatch dispersal in a gliding nymphal stick insect (*Extatosoma tiaratum*). J. Exp. Biol. 223, jeb226266.

Zolotov, V., Frantsevich, L. and Falk, E.-M. (1975). Kinematik der phototaktischen Drehung bei der Honigbiene *Apis mellifera* L. J. Comp. Physiol. A 97, 339–353.

