## Supplemental Material for "Walking in circles: Linking high- and low-level parameter scaling of visually guided and spontaneous turning behaviour"

### Supplementary Material

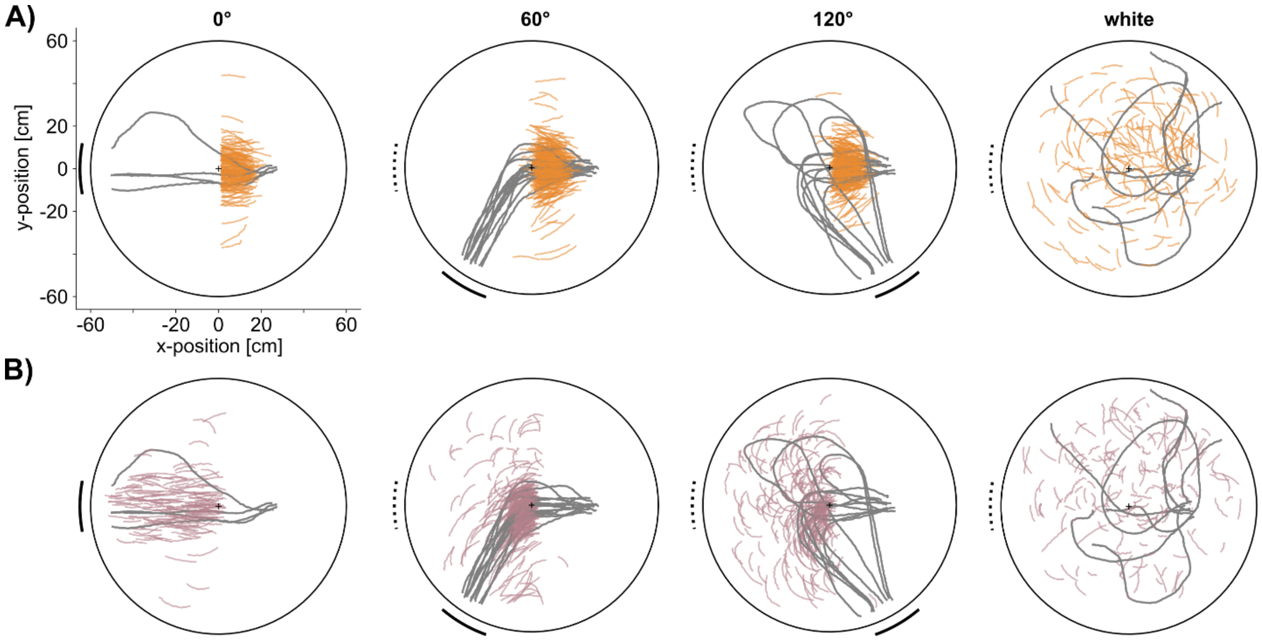

***Supplementary Figure 1:*** **Example trials and analysis episodes A and B.** Top view of the circular arena. Coloured trajectory parts show the prothorax position of '*on target*' trials. Each column corresponds to one of the four conditions, where 0°, 60° and 120° are "landmark trials" with (60°, 120°) and without displacement (0°) and "white" refers to an all-white reference condition without landmarks. Continuous underlying grey lines show exemplary trials of one randomly chosen animal. (**A**) In landmark trials, episode A started three seconds before the displacement of the black bar in 60° and 120° trials, or three seconds before crossing of the midline = control trials without displacement (0°). In all-white trials, it started half-way of the trial duration. (**B**) Episode B covered the three second period with largest mean rotation, after episode A.

**Supplementary Movie 1:** Example trial of animal walking through the arena in a 0° trial. Colored points show marker-positions obtained from DLC-tracking. Upper left white plot shows circular arena with black bar, prothorax position as well as front leg tibia-tarsus joint positions throughout the trial. Arrows show swing direction of right (cyan) and left (red) front legs respectively.

**Supplementary Movie 2:** Example trial of animal walking through the arena in a 60° trial. Colored points show marker-positions obtained from DLC-tracking. Upper left white plot shows circular arena with black bar, prothorax position as well as front leg tibia-tarsus joint positions throughout the trial. Arrows show swing direction of right (cyan) and left (red) front legs respectively.

**Supplementary Movie 3:** Example trial of animal walking through the arena in a 120° trial. Colored points show marker-positions obtained from DLC-tracking. Upper left white plot shows circular arena with black bar, prothorax position as well as front leg tibia-tarsus joint positions throughout the trial. Arrows show swing direction of right (cyan) and left (red) front legs respectively.

**Supplementary Movie 4:** Example trial of animal walking through the arena in an all-white trial. Colored points show marker-positions obtained from DLC-tracking. Upper left white plot shows circular arena with black bar, prothorax position as well as front leg tibia-tarsus joint positions throughout the trial. The Bar indicated in the beginning was not present but only indicates the starting orientation of the trial. Arrows show swing direction of right (cyan) and left (red) front legs respectively.

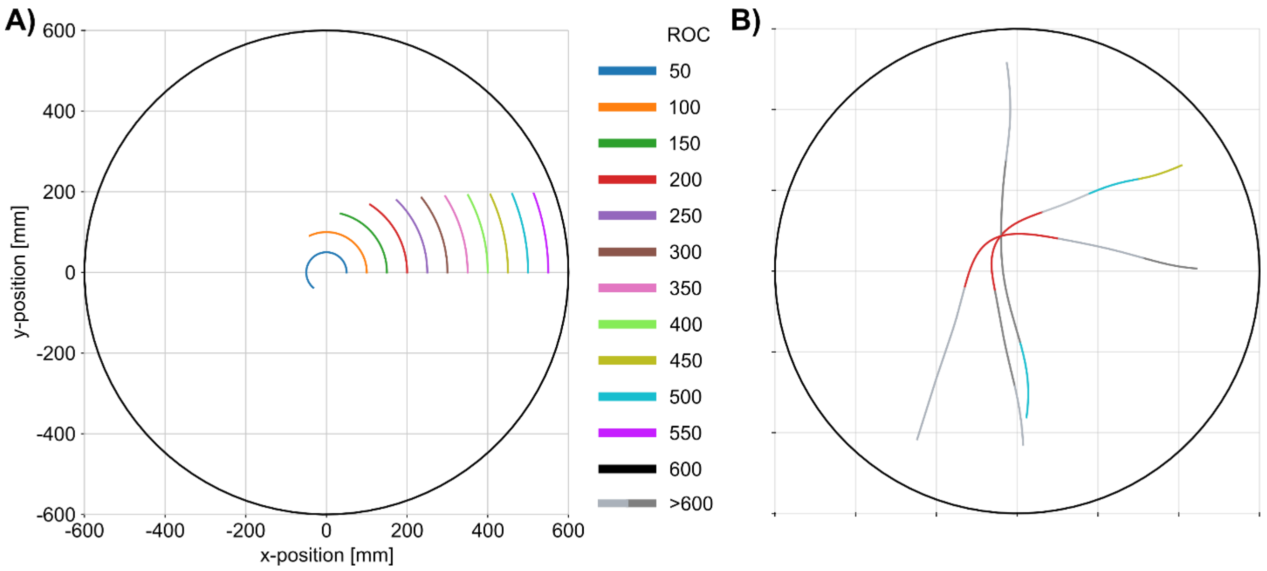

***Supplementary Figure 2: Trajectory segmentation according to curvature.*** (**A**) Schematic illustration of segments of the same length with different radius of curvature (ROC) in mm. The outer black circle shows the arena size (radius: 600 mm). (**B**) Segmented trajectories of three exemplary trials. Segments are coloured according to their ROC. Legend in (A) lists upper bounds of ROC ranges. For example, red segments have a ROC between 200 and 249 mm. Different grey segments have a ROC larger than 600 mm.

***Supplementary Table 1*****:** p-values of a priori post hoc tests on matched pairs (Wilcoxon's signed rank test) for 'at wall' and 'on target' trials.

| **Comparison** | **At wall** | **On target** |
|---|---|---|
| **Landmark vs. all-white** | <0.0001 | n.a. |
| **Starting orientation** | 0.17 | 0.92 |
| **Displacement vs. stationary** | 0.63 | 0.73 |
| **Displacement magnitude** | <0.0001 | <0.0001 |
| **Side of displacement** | 0.6 | 0.23 |

***Supplementary Table 2:*** Friedman test for matched samples results for differences in path length, curvature, median and maximum forward velocity. If Friedman test result was smaller than 0.05, Bonferroni-corrected Wilcoxon's signed rank post-hoc were done. $\alpha_{corr} = 0.05/6 = 0.0083$. Mean and standard error of the mean are given for each condition.

| | **Path length [mm]** | | **Curvature** | | **Median velocity [mm/s]** | | **Maximal velocity [mm/s]** | |
|---|---|---|---|---|---|---|---|---|
| **comparison** | Friedman | Wilcoxon | Friedman | Wilcoxon | Friedman | Wilcoxon | Friedman | Wilcoxon |
| | <0.0001 | | <0.0001 | | <0.0001 | | 0.36 | |
| **120 vs. 60** | | <0.0001 | | <0.0001 | | 0.006 | | |
| **120 vs. 0** | | <0.0001 | | <0.0001 | | <0.0001 | | |
| **120 vs. white** | | 0.015 | | <0.0001 | | <0.0001 | | |
| **0 vs. white** | | <0.0001 | | <0.0001 | | <0.0001 | | |
| **60 vs. 0** | | <0.0001 | | <0.0001 | | 0.347 | | |
| **60 vs. white** | | <0.0001 | | <0.0001 | | <0.0001 | | |
| | **mean [mm]** | **s.e.m. [mm]** | **mean** | **s.e.m.** | **mean [mm/s]** | **s.e.m. [mm/s]** | **mean [mm/s]** | **s.e.m. [mm/s]** |
| **0** | 987.33 | 28.81 | 1.31 | 0.04 | 39.66 | 1.17 | 142.64 | 2.73 |
| **60** | 1121.45 | 39.33 | 1.63 | 0.05 | 38.58 | 1.07 | 145.38 | 2.52 |
| **120** | 1465.30 | 36.75 | 3.07 | 0.09 | 34.67 | 1.07 | 143.48 | 2.89 |
| **white** | 1765.23 | 99.95 | 6.50 | 0.79 | 29.16 | 1.19 | 141.26 | 2.27 |

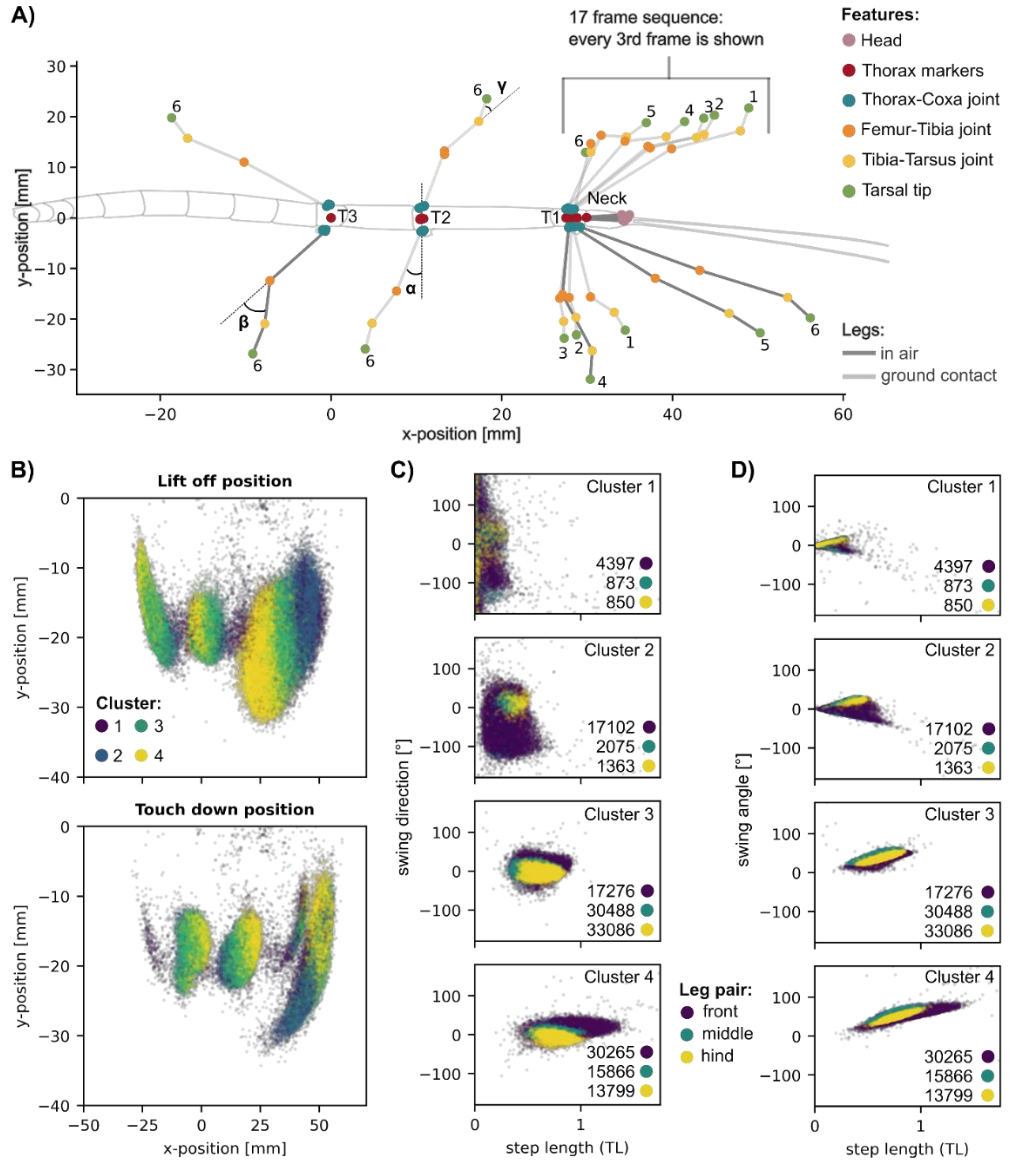

***Supplementary Figure 3:*** Classification of step cycle phase and type of step. (**A**) Classification of step cycle phase, distinguishing episodes "with ground contact" (stance) from episodes "with no ground contact" (swing). Schematic drawing of a stick insect with all tracked features, including exemplary results for stance in the left front leg and for swing in the right front leg. The three leg angles, α, β and γ are shown for right middle, right hind and left middle legs, respectively, but were calculated for all legs. Legs drawn in dark grey were classified as 'with no ground contact' (right front and hind legs), those drawn in light grey were classified as 'with ground contact'. For front legs, one leg every third frame was plotted from a temporal sequence of 17 frames. Numbers besides the legs indicate the temporal succession. (**B-D**) Step type classification. (**B**) Lift-off and touch-down position of four different clusters of steps for three leg pairs. Left legs were mirrored for plotting. Positions were centred on the metathorax marker T3. (**C**) Swing direction is plotted against step length for four different clusters; colours show different leg pairs. Numbers within each subplot show the number of steps per cluster and leg pair. (**D**) Swing angle is plotted against step length for four different clusters, with details as in C.

***Supplementary Table 3:*** Friedman tests for matched samples results for differences in percentage of three classes of steps between straight and curve walking and between inner and outer legs. If Friedman test result was smaller than 0.05, Bonferroni-corrected Wilcoxon’s signed rank post-hoc were done. $\alpha_{corr}$ = 0.05/3 = 0.0167. Mean and standard error of the mean are given for each condition.

| | **Inner leg** | **Outer leg** |
|---|---|---|
| **comparison** | Friedman<0.0001 | Friedman<0.0001 |
| **curve vs. straight, long steps** | $p<0.0001$ | $p<0.0001$ |
| **curve vs. straight, middle steps** | $p<0.0001$ | $p<0.0001$ |
| **curve vs. straight, short steps** | $p<0.0001$ | $p<0.0001$ |
| | **Straight** | **Curve** |
| **comparison** | Friedman<0.0001 | Friedman<0.0001 |
| **inner vs. outer, long steps** | $p=0.0003$ | $p<0.0001$ |
| **inner vs. outer, middle steps** | $p=0.0006$ | $p<0.0001$ |
| **inner vs. outer, short steps** | $p=0.44$ | $p=0.03$ |

| | **Inner leg** | | **Outer leg** | | |
|---|---|---|---|---|---|
| | **mean [%]** | **s.e.m. [%]** | | **mean [%]** | **s.e.m. [%]** |
| **curve long** | 59.42 | ± 2.32 | **curve long** | 77.98 | ± 1.78 |
| **curve middle** | 33.51 | ± 1.99 | **curve middle** | 16.05 | ± 1.36 |
| **curve short** | 7.07 | ± 1.3 | **curve short** | 5.97 | ± 0.84 |
| **straight long** | 74.56 | ± 1.84 | **straight long** | 77.22 | ± 1.74 |
| **straight middle** | 20.45 | ± 1.52 | **straight middle** | 17.84 | ± 1.53 |
| **straight short** | 4.99 | ± 0.9 | **straight short** | 4.94 | ± 0.84 |

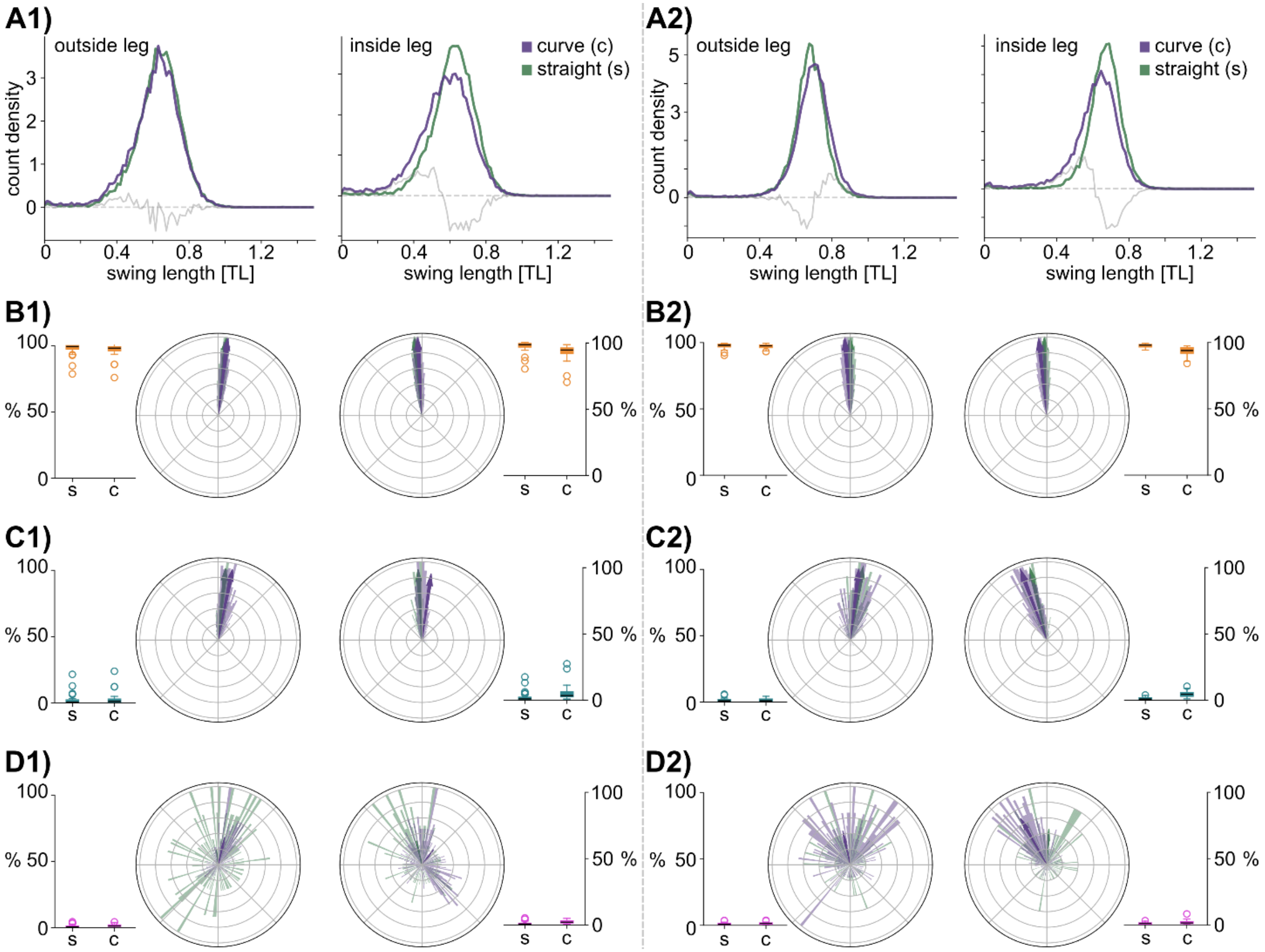

***Supplementary Figure 4: Three classes of step types in middle and hind legs**. Same graph details as in Fig. 7 of main manuscript, but for different leg types. (**A**) Density plots showing the distribution of swing lengths of middle (1) and hind (2) legs (left: outer leg, right: inner leg) in curved (purple, ROC <= 600, N=32, Middle legs: $n_{steps\_inside}$= 15706, $n_{steps\_outside}$= 16919; Hind legs: $n_{steps\_inside}$= 15570, $n_{steps\_outside}$= 16812) and straight segments (turquoise, ROC > 600, N=32, Middle legs: $n_{steps\_inside}$ = 12200, $n_{steps\_outside}$ = 12263; Hind legs: $n_{steps\_inside}$ = 12190, $n_{steps\_outside}$ = 12291). The grey solid line shows the difference between the curved and straight condition. The grey dashed line highlights zero density. N = 32 (**B-D**) Circular histograms showing the swing direction of steps of front legs (left column: outside leg, right column: inside leg) in curved (purple) and straight (green) segments. The mean direction (dir) and the concentration around mean direction (r) are given below. Boxplots show the percentage of a step type per condition as means per animal. N = 31. (**B**) Long steps. (**C**) Middle steps. (**D**) Short steps.*

***Supplementary Table 4:*** Friedman test for matched samples results for differences in overall rotation and step type asymmetry. If Friedman test result was smaller than 0.05, Bonferroni-corrected Wilcoxon's signed rank post-hoc were done. $\alpha_{corr}$ = 0.05/4 = 0.0125. Mean and standard error of the mean are given for each condition.

| | **Rotation [°]** | | **Asymmetry** | |
|---|---|---|---|---|
| | Friedman p<0.0001 | | Friedman p<0.0001 | |
| **60 e. A vs. 120 e. A** | p = 0.9 | | p = 0.02 | |
| **60 e. C vs.120 e. C** | p<0.0001 | | p = 0.0015 | |
| **60 e. A vs. 60 e. C** | p<0.0001 | | p<0.0001 | |
| **120 e. A vs. 120 e. C** | p<0.0001 | | p<0.0001 | |
| | **mean [°]** | **s.e.m. [°]** | **mean** | **s.e.m.** |
| **60 e. A** | 11.42 | 0.81 | 0.99 | 0 |
| **60 e. C** | 57.56 | 0.98 | 0.88 | 0.02 |
| **120 e. A** | 11.4 | 1.2 | 0.97 | 0.01 |
| **120 e. C** | 88.28 | 2.46 | 0.73 | 0.04 |

***Supplementary Table 5:*** Friedman test for matched samples results for differences in change of swing direction of long steps between episode A and B. If Friedman test result was smaller than 0.05, Bonferroni-corrected Wilcoxon's signed rank post-hoc were done. $\alpha_{corr}$ = 0.05/4 = 0.0125. Mean and standard error of the mean are given for each condition.

| | **Direction** | | **Variance** | |
|---|---|---|---|---|
| | Friedman p<0.0001 | | Friedman p<0.0001 | |
| **60 inside vs. 120 inside** | p=0.0048 | | p=0.2941 | |
| **60 outside vs.120 outside** | p=0.0001 | | p=0.0534 | |
| **60 inside vs. 60 outside** | p=0.3579 | | p<0.0001 | |
| **120 inside vs. 120 outside** | p=0.7172 | | p<0.0001 | |
| | **mean** | **s.e.m.** | **mean** | **s.e.m.** |
| **60 inside** | 7.07 | 0.63 | 0.0243 | 0.0033 |
| **60 outside** | 7.91 | 0.61 | -0.0007 | 0.0014 |
| **120 inside** | 14.7 | 2.62 | 0.0226 | 0.0071 |
| **120 outside** | 14.38 | 1.24 | -0.0062 | 0.0022 |

***Supplementary Table 6:*** Friedman test for matched samples results for differences in change of swing direction of middle steps between episode A and B. Mean and standard error of the mean are given for each condition below.

| | **Direction** | | **Variance** | |
|---|---|---|---|---|
| | Friedman p=0.05 | | Friedman p=0.22 | |
| | **mean** | **s.e.m.** | **mean** | **s.e.m.** |
| **60 inside** | 30.17 | 9.45 | 0.0881 | 0.0411 |
| **60 outside** | -17.54 | 14.59 | -0.0137 | 0.0627 |
| **120 inside** | 28.43 | 12.44 | 0.1401 | 0.0615 |
| **120 outside** | 2.19 | 10.87 | 0.0858 | 0.0868 |

***Supplementary Table 7:*** Friedman test for matched samples results for differences in percentages of short steps for different ROCs. Bonferroni-corrected Wilcoxon's signed rank tests were done post-hoc. $\alpha_{corr}$ = 0.05/4 = 0.0125. Mean and standard error of the mean are given for each condition on the left.

| | **mean [%]** | **s.e.m. [%]** | | Friedman p<0.0001 |
|---|---|---|---|---|
| **1-200 white** | 6.08 | 0.73 | | **Wilcoxon**: |
| **201-400 white** | 3.79 | 0.94 | **white vs stim. (1-200)** | p=0.0024 |
| **401-600 white** | 4.12 | 0.77 | **white vs stim. (201-400)** | p=0.9205 |
| **601-4000 white** | 3.84 | 0.66 | **white vs stim. (401-600)** | p=0.7835 |
| **1-200 stim** | 4.6 | 0.57 | **white vs stim. (601-4000)** | p=0.0385 |
| **201-400 stim** | 3.48 | 0.55 | | |
| **401-600 stim** | 3.49 | 0.57 | | |
| **601-4000 stim** | 2.73 | 0.42 | | |
